# Cdc48 influence on separase levels is independent of mitosis and suggests translational sensitivity of separase

**DOI:** 10.1101/2021.04.28.441771

**Authors:** Drisya Vijayakumari, Janina Müller, Silke Hauf

## Abstract

Cdc48 (p97/VCP) is a AAA-ATPase that can extract ubiquitinated proteins from their binding partners and channel substrates to the proteasome. A fission yeast *cdc48* mutant (*cdc48-353*) shows low levels of the cohesin protease, separase, and pronounced chromosome segregation defects in mitosis. Separase initiates chromosome segregation when its binding partner securin is ubiquitinated and degraded. The low separase levels in the *cdc48-353* mutant have been attributed to a failure to extract ubiquitinated securin from separase resulting in co-degradation of separase along with securin. If true, this establishes Cdc48 as a key regulator of mitosis. In contrast, we show here that low separase levels in the *cdc48-353* mutant are independent of mitosis. Moreover, we find no evidence of enhanced separase degradation in the mutant. Instead, we suggest that the *cdc48-353* mutant uncovers specific requirements for separase translation. Our results highlight a need to better understand how this key mitotic enzyme is synthesized.

## Introduction

Separase is a protease that triggers chromosome separation in anaphase by cleaving the cohesin complex (Uhlmann et al., 2000). Prior to anaphase, separase is kept inactive by inhibitory binding of securin, as well as in some eukaryotes by inhibitory binding of cyclin B and Sgo2/Mad2 (Hellmuth et al., 2020; Stemmann et al., 2001; Uhlmann, 2001). Removal of securin from separase is initiated by the anaphase promoting complex (APC/C), an E3 ligase. The APC/C ubiquitinates securin, which marks it for proteasomal degradation. The APC/C also ubiquitinates cyclin B, whose degradation then leads to the inactivation of cyclin-dependent kinase (CDK1) and mitotic exit (Peters, 2006).

Cdc48 (also known as p97 or VCP in vertebrates) is a homohexameric AAA-ATPase that targets ubiquitinated proteins and is essential for cell viability. Cdc48 can unfold ubiquitinated proteins and promote their degradation by the proteasome (van den Boom and Meyer, 2018; Ye et al., 2017). Cdc48 has an N-terminal domain, two tandem ATPase domains, D1 and D2, and an unstructured C-terminal tail. In the hexameric complex, D1 and D2 form a double ring structure with a central pore through which substrates can be threaded, which unfolds the polypeptide (Banerjee et al., 2016; Bodnar and Rapoport, 2017; Cooney et al., 2019; Twomey et al., 2019). Both the N-terminal domain and the C-terminal tail provide platforms for cofactor binding. Two of the best studied cofactors are the heterodimeric Ufd1/Npl4 complex and Shp1 (p47), which mediate substrate recognition (Buchberger et al., 2015; Schuberth et al., 2004; Ye et al., 2017). Other cofactors change the degree or type of ubiquitination of substrates (Jentsch and Rumpf, 2007; Ye et al., 2017). Ufd2 is one such cofactor, which in yeast binds to the C-terminal tail of Cdc48 and acts as E4 ubiquitin ligase, adding additional ubiquitin moieties to enhance proteasomal degradation (Böhm et al., 2011; Rumpf and Jentsch, 2006).

Central to the function of Cdc48 is its role as a segregase. Cdc48 separates ubiquitinated targets from their non-modified binding partners or from cellular structures (Stolz et al., 2011; Ye et al., 2017). For example, p97^Ufd1/Npl4^ extracts ubiquitinated Aurora B kinase from chromatin at mitotic exit to promote chromatin decondensation and nuclear envelope reformation (Ramadan et al., 2007). In a similar way, yeast Cdc48 mobilizes the transcription factors Mga2 and Spt23 from the ER membrane (Rape et al., 2001; Shcherbik and Haines, 2007).

In fission yeast (*S. pombe*), a temperature sensitive allele of *cdc48* (*cdc48-353*) was isolated based on its characteristic of being suppressed by separase overexpression (Yuasa et al., 2004). The mutation in *cdc48-353* resides in the D1 domain and replaces glycine 338 with aspartate (G338D) (Ikai and Yanagida, 2006). The *cdc48-353* mutant strain has low levels of separase and displays severe chromosome segregation defects, similar to separase mutants. This segregation defect is rescued by overexpression of separase (Ikai and Yanagida, 2006; Yuasa et al., 2004). To explain this phenotype, it has been proposed that Cdc48 may target polyubiquitinated securin and extract it from separase (Yuasa et al., 2004). The low separase levels in the *cdc48-353* mutant could then be explained by a failure to extract securin from separase and subsequent codegradation of separase along with securin during anaphase (Fig. 1A). This is an attractive model that is consistent with findings that impairing Cdc48/p97 activity can lead to mitotic delays in budding yeast and human cells (Chien and Chen, 2013; Wojcik et al., 2004), and that the Cdc48 ortholog p97 recognizes branched ubiquitin chains synthesized by the human APC/C (Meyer and Rape, 2014; Oh et al., 2020). If the hypothesis is true, Cdc48 would be a key player in the regulation of anaphase.

**Figure 1.**
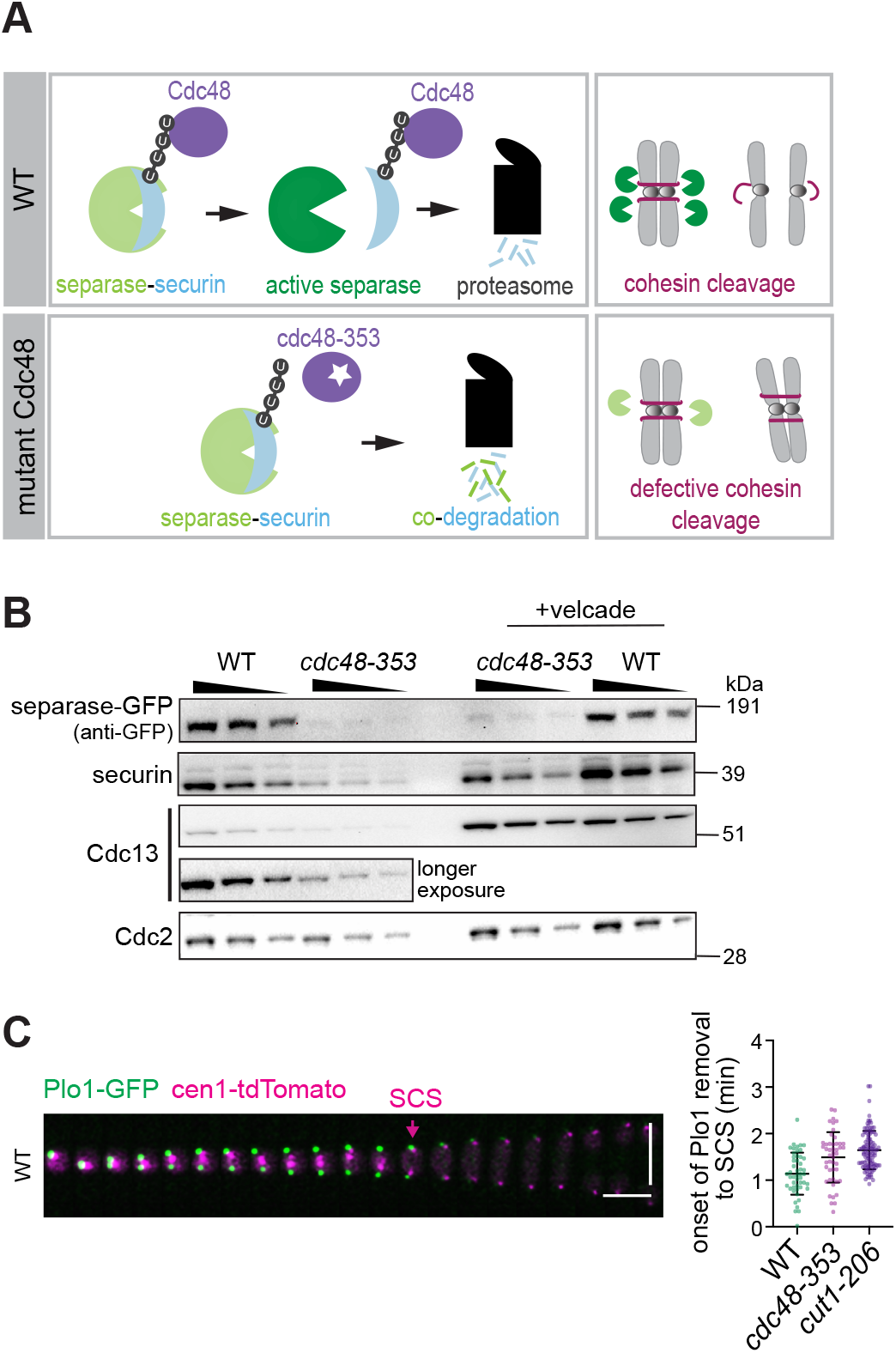
Mitotic phenotype of *cdc48-353* cells and previously proposed model. **(A)** Schematic depiction of the model which proposes failure of Cdc48 segregase activity in the *cdc48-353* mutant, leading to co-degradation of separase and securin during mitosis. **(B)** Immunoblot showing separase(Cut1)-GFP, securin (Cut2) and Cdc13 (cyclin B) levels in the wild type (WT) or *cdc48-353* mutant with and without 45 min treatment with the proteasome inhibitor velcade. A 1:1 dilution series is loaded for each sample to allow for semi-quantitative comparison. Cdc2 serves as loading control. **(C)** Left: Representative kymograph of a WT cell during mitosis. Cells were expressing Plo1-GFP as marker for mitotic entry and exit, and a centromeric tdTomato marker to analyze sister chromatid separation (SCS). Vertical scale bar: 10 μM, horizontal scale bar: 2 min. Right: Comparison of WT, *cdc48-353* and *cut1-206* strains for the time taken from onset of Plo1 removal from spindle pole bodies (decline in CDK1 activity) to SCS. Black bars are mean and standard deviation.

Here, we have tested—and falsify—this hypothesis. Using a combination of live cell imaging, genetics and biochemistry in fission yeast (*S. pombe*), we demonstrate that separase is not specifically degraded during mitosis in the *cdc48-353* mutant. Moreover, we generally do not find evidence of enhanced separase degradation. Instead, we argue that the *cdc48-353* mutant uncovers that separase has specific requirements for its translation, which is an aspect of separase regulation that has been little studied.

## Results

### Securin mutants that lack the ability to bind separase are not toxic when overexpressed in the *S. pombe cdc48-353* mutant

The levels of separase and those of the key APC/C substrates securin and Cdc13 (the *S. pombe* mitotic cyclin B) are reduced in the *cdc48-353* mutant (Fig. 1B) (Ikai and Yanagida, 2006). As we will discuss in more detail below, treatment with the proteasome inhibitor Velcade (bortezomib) did not restore separase levels in the *cdc48-353* mutant, although it increased securin and Cdc13 levels (Fig. 1B). Consistent with the low levels of separase, we found that sister chromatid separation was delayed in *cdc48-353* mutant cells relative to the decline in CDK1 activity at mitotic exit (Fig. 1C). This strongly resembles the defects seen in a separase mutant (*cut1-206*) and thus suggests that the mitotic phenotype of the *cdc48-353* mutant is caused by reduced separase activity.

Consistent with low separase activity in *cdc48-353* mutant cells, overexpression of the separase inhibitor securin becomes lethal (Ikai and Yanagida, 2006; Yuasa et al., 2004). We found this to be the case even if securin levels were elevated to only about 8-times the wild-type level (Fig. 2A). To better understand this toxicity, we screened for securin mutants whose overexpression is tolerated in *cdc48-353* mutant cells (Fig. 2B). Of ten such mutants that we sequenced, six were truncations of securin that terminated before the separase binding motif (SBM) (Boland et al., 2017; Lin et al., 2016; Luo and Tong, 2017; Nagao and Yanagida, 2006). The other four mutants had missense or frameshift mutations clustered around the SBM. The SBM of securin is known to bind into the separase catalytic site (Boland et al., 2017; Lin et al., 2016; Luo and Tong, 2017; Nagao and Yanagida, 2006). Overexpression of the previously characterized *cut2AIA* mutant (DIE in the SBM replaced with AIA; (Nagao et al., 2004)) or of a mutant that lacks the SBM and all sequences downstream (*cut2N121*), was also tolerated in the *cdc48-353* mutant background. In contrast, the expression of wild-type securin (*cut2^+^*) from the same promoter was lethal (Fig. 2C). Hence, the SBM is crucial for the toxic effect. The levels of all overexpressed securin versions (both mutant and wild-type) were drastically lowered in the *cdc48-353* mutant background, suggesting that the mutants remain susceptible to the effects of the *cdc48-353* mutant (Fig. 2D).

**Figure 2.**
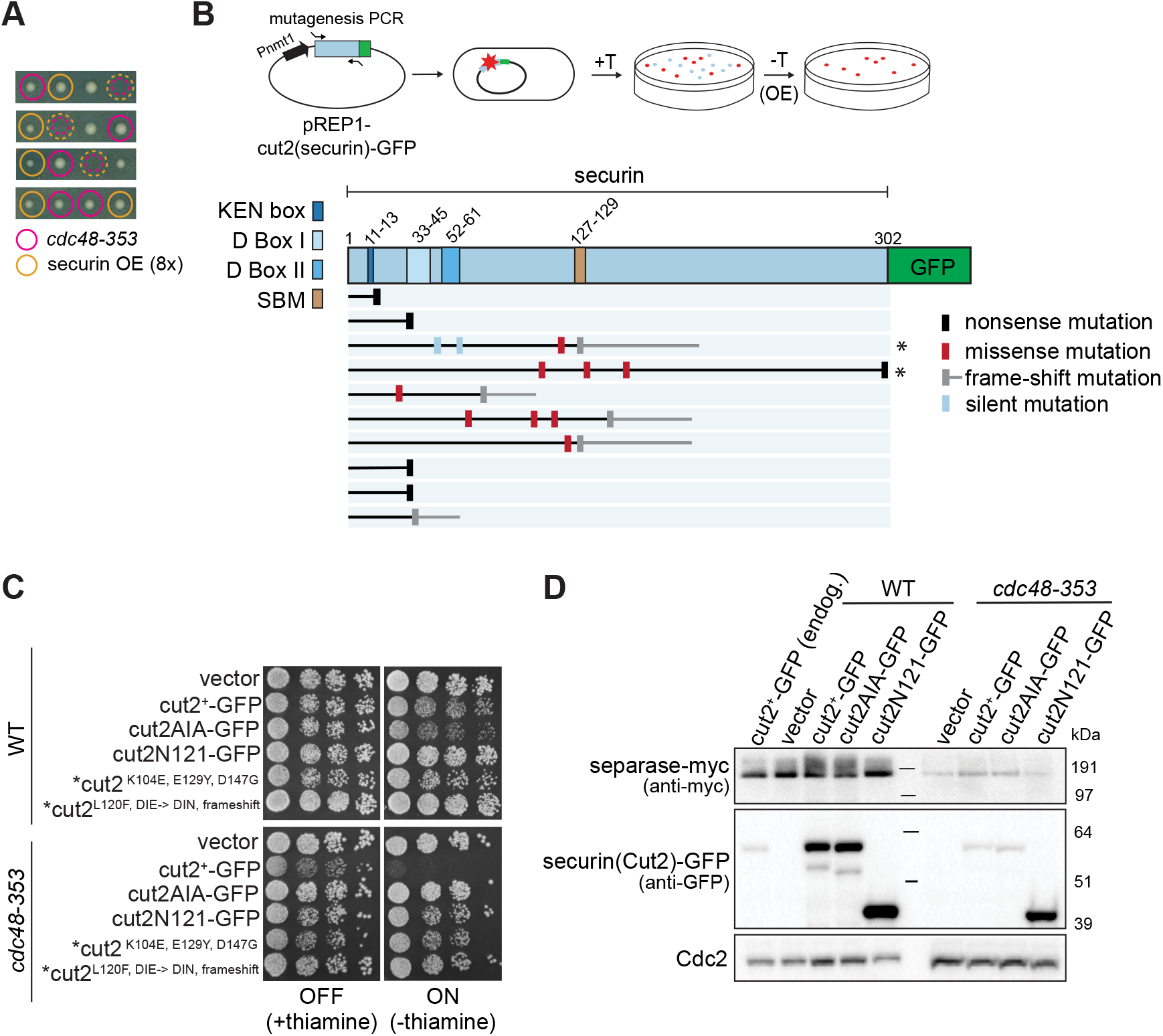
Overexpression of securin mutants compromised for interaction with separase is tolerated in the *cdc48-353* mutant. **(A)** Tetrads from a cross between *cdc48-353* mutant and a strain overexpressing (OE) securin to about 8-times the wild-type level. Colored circles indicate the strain genotype, dashed colored circles indicate spore lethality. **(B)** Top: Schematic of securin mutagenesis screen to identify mutations that lose toxicitiy in *cdc48-353* cells upon overexpression. Securin expression from the plasmid is suppressed in the presence of thiamine (+T) and induced in its absence (-T). Mutations that have lost toxicity when overexpressed are indicated in red. Bottom: Schematic summary indicating positions and type of mutations identified in the non-toxic mutants. The length of the protein formed in each case is indicated with a black or grey horizontal line. Mutants examined in (C) are marked by an asterisk. SBM = separase-binding motif (aka SIS = separase interaction segment). **(C)** Growth assay of cells inducibly expressing the indicated securin (*cut2*) mutants from the *nmt81* promoter in the wild-type or *cdc48-353* mutant background. Cells were grown on minimal medium in the presence or absence of thiamine. The latter induces expression. The strains indicated with asterisks are those retrieved directly from the screen. **(D)** Immunoblot of separase and securin levels in the wild type and *cdc48-353* mutant overexpressing the indicated variants of securin (*cut2*) or an empty vector, 15 hours after induction. A strain expressing securin-GFP from the endogenous locus is included as reference in the first lane. Cdc2 serves as loading control.

The securin mutants that lacked toxicity when overexpressed presumably neither bind nor inhibit separase efficiently (Nagao et al., 2004; Nagao and Yanagida, 2006). Hence, these mutants may lack toxicity because they are unable to mediate separase co-degradation with securin due to their failure to bind separase, or they may lack toxicity because they do not inhibit the small, remaining amount of separase in the *cdc48-353* mutant. When we compared short-term overexpression of the toxic wild-type securin (Cut2) and the non-toxic Cut2AIA in *cdc48-353* mutant cells, we did not observe any difference in the amount of separase (Fig. 2D). These two mutants should have been separase-binding and non-binding, respectively, and the result therefore indicates that overexpression of wild-type securin does not further enhance separase degradation. This casts doubt on whether separase indeed undergoes co-degradation with securin in *cdc48-353* mutant cells.

### Separase levels do not drop during mitosis in the *cdc48-353* mutant

To directly test the proposed model of co-degradation (Fig. 1A), we imaged cells expressing securin-GFP or separase-GFP as they underwent mitosis. If Cdc48 indeed aided the proteasome-mediated degradation of securin during mitosis, securin might be expected to degrade slower in the *cdc48-353* mutant. The level of securin-GFP in the *cdc48-353* mutant was lower than in the wild-type background (Fig. 3A; S1A), consistent with the finding for endogenous securin by immunoblot (Fig. 1B). However, the cellular degradation kinetics of securin-GFP were indistinguishable in *cdc48^+^* and *cdc48-353* mutant cells after normalizing for the reduced level (Fig. 3A). Similar results were obtained for Cdc13-GFP (Fig. S1C,D). Hence, securin and Cdc13 are still efficiently targeted for proteasomal degradation in the *cdc48-353* mutant.

**Figure 3.**
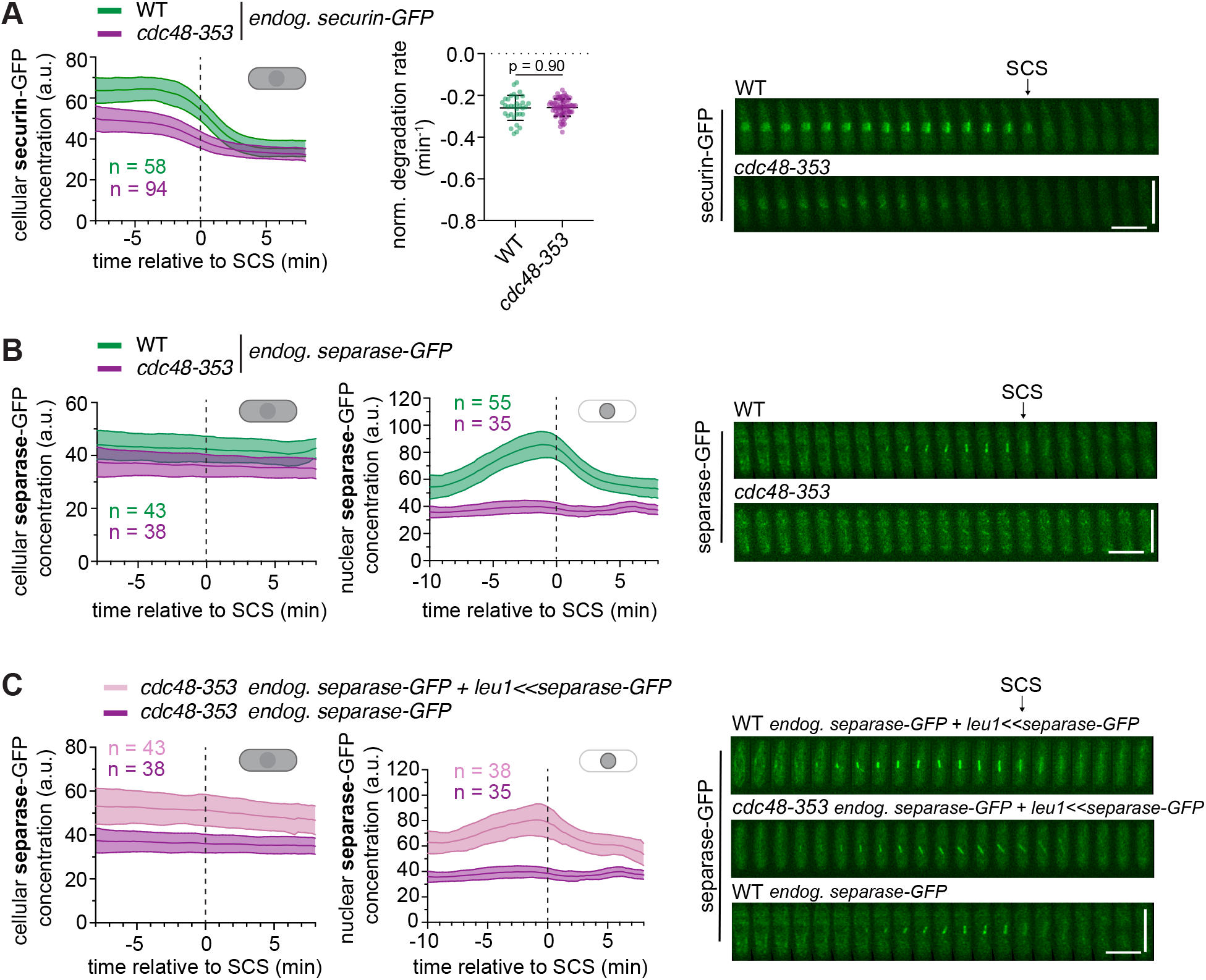
Separase levels in the *cdc48-353* mutant do not drop during mitosis. **(A)** Left: Cellular concentrations of securin-GFP in the wild type (WT) and *cdc48-353* mutant during mitosis. Mean (central line) and standard deviation (area) are shown (n = number of cells). Data are aligned to the time of sister chromatid separation (SCS), indicated by the vertical dashed line. Middle: The normalized degradation rate of cellular securin during mitosis. Bars are mean and standard deviation. Statistical significance was tested by unpaired t-test. Right: Representative kymographs showing securin-GFP in the WT and *cdc48-353* mutant as cells undergo mitosis. The point of SCS is indicated by an arrow. Horizontal scale bar: 2 min, vertical scale bar: 10 μM. **(B)** The cellular (left) and nuclear (middle) concentrations of separase-GFP in the WT and *cdc48-353* mutant during mitosis. Right: Representative kymographs showing separase-GFP in the WT and *cdc48-353* mutant as cells progress through mitosis. **(C)** The cellular (left) and nuclear (middle) concentrations of separase-GFP in the *cdc48-353* mutant, either expressing separase-GFP only from the endogenous locus, or additionally expressing a second copy of separase-GFP from the exogenous *leu1* locus. The data for the strain with only endogenous separase-GFP are the same as in (B). Right: Kymographs showing separase-GFP in the WT and *cdc48-353* mutant. The cells in the top two kymographs were expressing a second copy of separase-GFP from the exogenous *leu1* locus.

We next checked if a decline in separase levels occurs during mitosis. Similar to securin-GFP, both the cellular and nuclear levels of separase-GFP were low in *cdc48-353* mutant cells (Fig. 3B; S1B). However, we did not observe a drop in separase-GFP levels during mitosis as would be expected if separase was co-degraded with securin (Fig. 3B). Given the already low separase levels of the *cdc48-353* mutant, a further drop may be difficult to detect. To circumvent this issue, we expressed a second copy of separase-GFP under its endogenous promoter from another genetic locus (*leu1*). A second copy of separase was sufficient to rescue growth of the *cdc48-353* mutant cells, but only if it was catalytically active (Fig. S2A). The cellular and nuclear separase levels now were indeed increased, but we were still unable to observe any drop during mitosis (Fig. 3C; S2B).

To separately assess the effect of *cdc48-353* on the two copies of separase, we used two different tags: GFP at the endogenous and myc at the exogenous locus. Separase-myc expressed from the exogenous locus remained susceptible to the *cdc48-353* mutant as its levels were also lowered (Fig. S2D). The presence of this extra copy did not change the level of separase-GFP expressed from the endogenous locus (Fig. S2C,D). Hence, the mechanism that lowers separase levels in the *cdc48-353* mutant acts equally on separase expressed from the endogenous and the exogenous locus. Securin levels were moderately higher in *cdc48-353* mutant cells with two copies of separase compared to one copy (Fig. S2D), consistent with findings that separase is able to stabilize securin (Kumada et al., 1998).

To further test the original model of securin and separase co-degradation, we anchored Cdc48 away from the nucleus (Fig. S2E) (Ding et al., 2014; Haruki et al., 2008). Despite successfully reducing the amount of Cdc48 in the nucleus (Fig. S2E), we did not observe an obvious decline of separase levels during mitosis (Fig. S2F).

Taken together, the unaltered securin degradation kinetics and stable separase levels throughout mitosis in the *cdc48-353* mutant suggest that the reduction in separase levels does not occur as a consequence of mitotic co-degradation with securin. Supporting this notion, we did not find a convincing physical association between Cdc48 and separase in asynchronous cultures (Fig. S3A), or between securin and Cdc48 in cells undergoing mitosis (Fig. S3B).

### *Cdc48-353* is a dominant negative gain-of-function allele

The *cdc48-353* allele causes a single amino acid change (G338D) in the D1 domain of Cdc48 within a loop (loop 2) that faces the central pore (Ikai and Yanagida, 2006; Marinova et al., 2015). In the D2 domain, pore-facing loops contain aromatic residues that are used to translocate substrates (Twomey et al., 2019). Pore-facing D1 loops of archaeal Cdc48 homologs retain aromatic residues, but in eukaryotic Cdc48 they have been replaced by non-aromatic residues (Esaki et al., 2017; Twomey et al., 2019). In budding yeast (*S. cerevisiae*), re-inserting an aromatic residue into the pore-facing loop 1 of D1 (e.g. M288Y) led to lethality (Esaki et al., 2017). Overexpression of the *S. cerevisiae* Cdc48M288Y mutant in the wild-type background was lethal as well, and this lethality was rescued by deletion of some, but not all, Cdc48 cofactors, by truncation of the Cdc48 C-terminal tail, or by impairing the ATPase activity of the D1 domain (Esaki et al., 2017). Interestingly, the *S. pombe cdc48-353* mutant has also been shown to be rescued by deletion of the cofactor Ufd2, by a specific mutation in the cofactor Ufd1, by mutations in the Cdc48 C-terminal tail, or by mutations expected to lower the D1 ATPase activity (Marinova et al., 2015; Xu et al., 2018). This suggested that the mechanism behind the toxicity of *S. pombe* Cdc48G338D and *S. cerevisiae* Cdc48M288Y may be similar.

It was previously reported that *S. pombe* strains overexpressing Cdc48G338D retained viability (Ikai and Yanagida, 2006). In our hands, though, *nmt1* promoter-driven overexpression of Cdc48G338D, but not wild-type Cdc48, strongly reduced separase and securin levels, caused abnormal cell division and was lethal (Fig. 4; S4). These phenotypes were rescued to a large extent by introducing an additional E325A mutation in the Walker B motif of the D1 domain (Fig. 4; S4B), which is expected to lower ATPase activity (Bodnar and Rapoport, 2017). Since C-terminal tagging of Cdc48-353 ameliorates the mutant phenotype (Marinova et al., 2015), these constructs were N-terminally tagged to follow their expression. The difference to previous results (Ikai and Yanagida, 2006) may be due to different levels of overexpression or different tagging of Cdc48. Based on our results, we conclude that Cdc48G338D behaves as dominant-negative.

**Figure 4.**
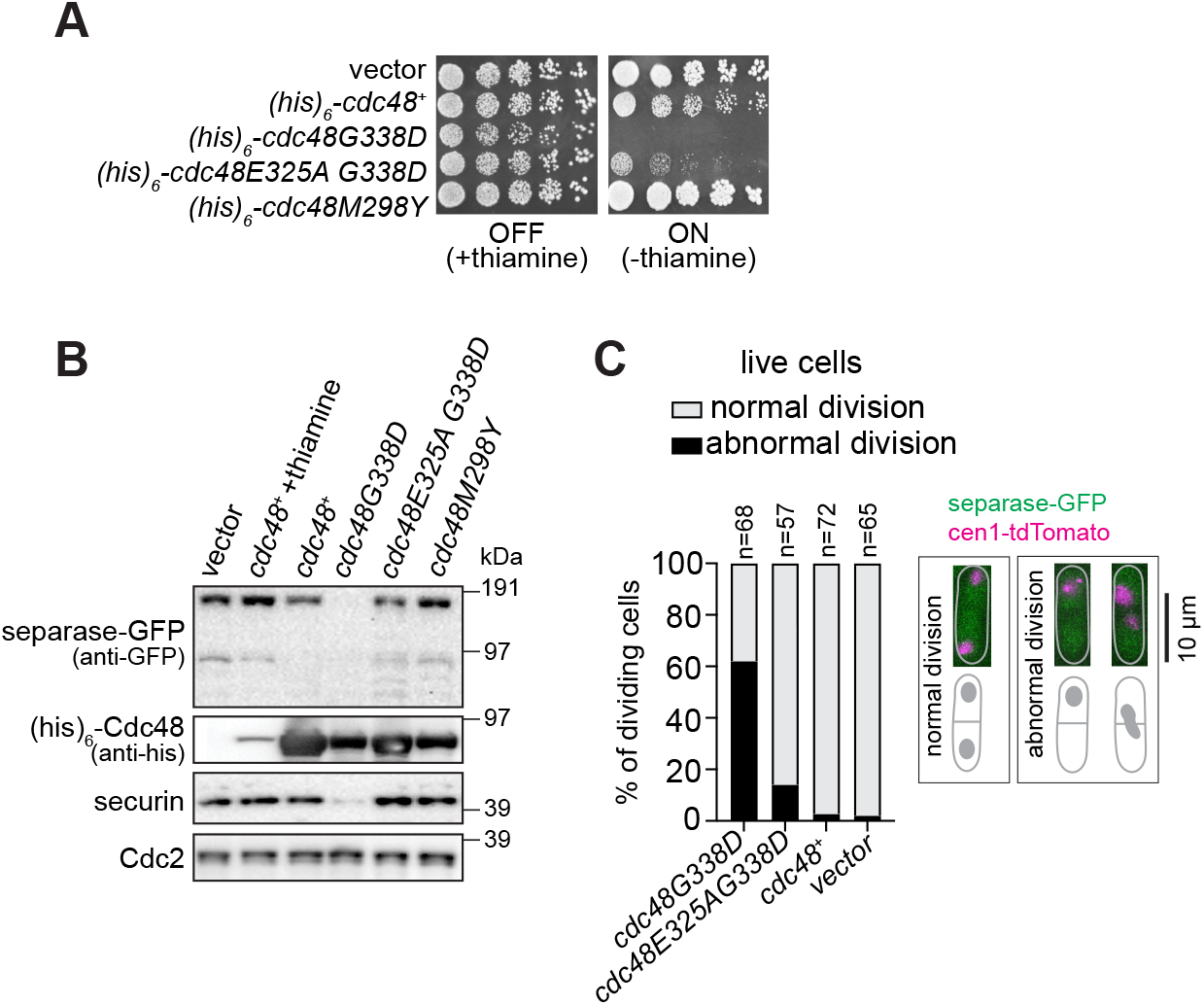
*Cdc48-353* is a dominant negative gain-of-function allele. **(A)** Growth of wild-type strains transformed with vectors for conditional overexpression of wild-type *cdc48^+^, cdc48* mutants, or the empty vector. Growth on minimal medium at 30°C in the presence or absence of thiamine; the latter induces expression. **(B)** Immunoblot showing separase-GFP and securin levels in strains overexpressing wild-type or mutant *cdc48*. Cdc48 levels probed with anti-his antibody show the level of Cdc48 overexpression in each strain (growth in thiamine-depleted conditions for 15 hrs, except for lane 2, for which cells were grown in the presence of thiamine). Cdc2 serves as loading control. **(C)** Quantification of cellular phenotypes in dividing cells overexpressing the indicated Cdc48 protein. Abnormal division includes cells with cut phenotype, unequal nuclear division or chromosome bridges.

We also generated the mutant analogous to *S. cerevisiae* M288Y, *S. pombe* Cdc48M298Y. However, overexpression of this mutant neither lowered separase levels nor was lethal (Fig. 4A,B). Furthermore, some cofactor deletions (shp1/ubx3Δ or ubx4Δ) which rescued the lethality of Cdc48M288Y overexpression in *S. cerevisiae* did not rescue *S. pombe cdc48-353* mutant growth (Fig. S5A). However, both *S. cerevisiae* Cdc48M288Y overexpression and *S. pombe cdc48-353* are rescued by *ufd2* deletion (Fig. 5A) (Esaki et al., 2017; Xu et al., 2018). Altogether, this indicates that G338D and M298Y do not cause the same cellular phenotype, although it remains possible that they mechanistically cause similar defects in Cdc48 function. For example, both may cause misprocessing of Cdc48 substrates, but possibly of different substrates.

**Figure 5.**
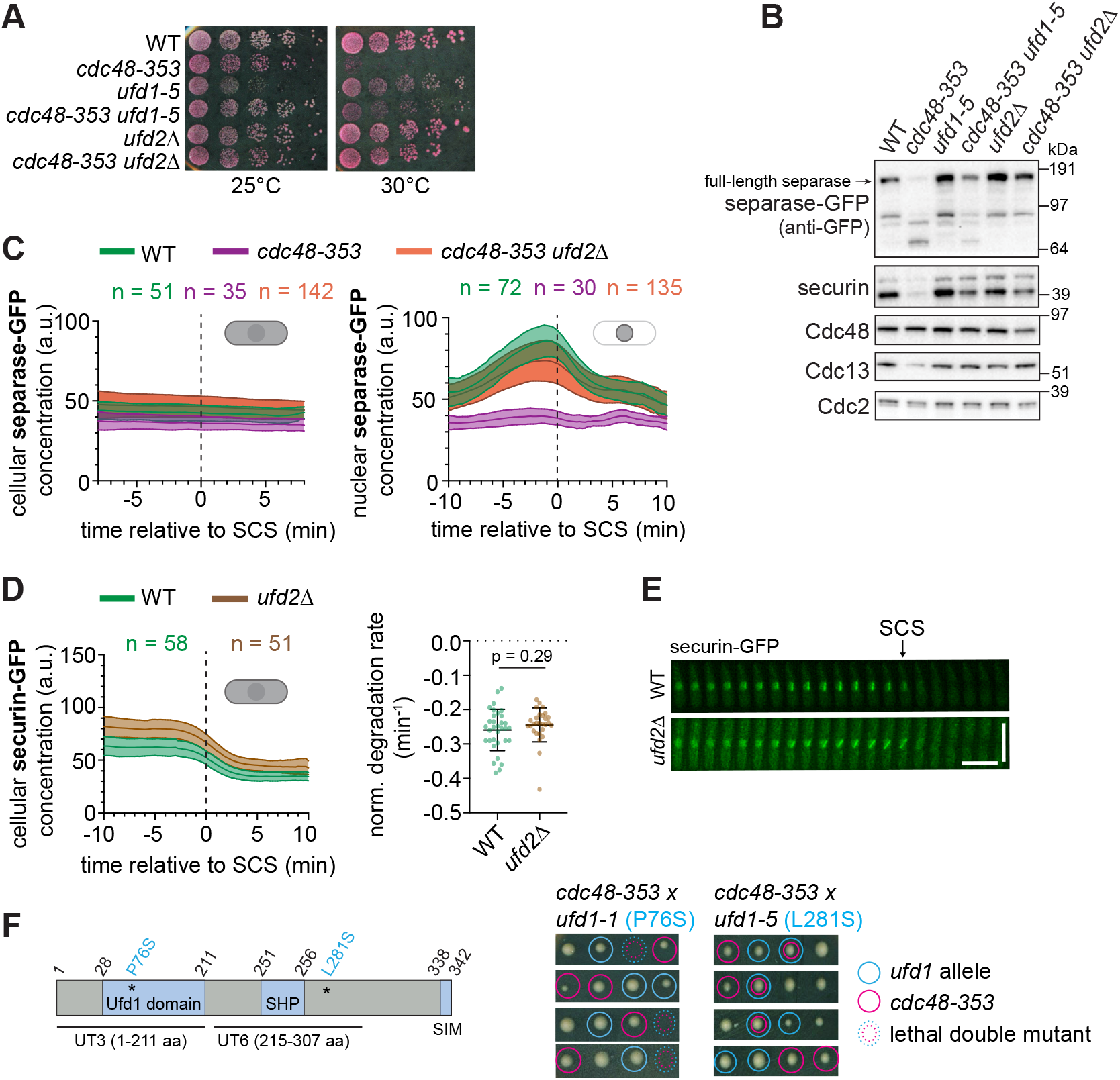
Compromised interaction with the cofactors Ufd1 and Ufd2 rescues separase levels in the *cdc48-353* mutant. **(A)** Growth assay of wild-type (WT) and the indicated single and double mutant strains at different temperatures on rich medium with Phloxine B, which stains dead cells. **(B)** Immunoblot showing the levels of the indicated proteins in the WT, *cdc48-353, ufd1-5* and *ufd2Δ* single and double mutants. **(C)** The cellular (left) and nuclear (right) concentrations of separase-GFP in the WT, *cdc48-353* single mutant, and *cdc48-353 ufd2Δ* double mutant during mitosis. Mean (central line) and standard deviation (area) are shown (n = number of cells). Data are aligned to the time of sister chromatid separation (SCS), indicated by the vertical dashed line. **(D)** Left: The cellular concentration of securin-GFP in the WT and *ufd2Δ* mutant during mitosis. Right: The normalized degradation rate of cellular securin during mitosis. Bars are mean and standard deviation. Statistical significance was tested by unpaired t-test. The WT data are the same as in Figure 3. **(E)** Kymographs showing securin-GFP in the WT and *ufd2Δ* mutant as cells progress through mitosis. Horizontal scale bar: 2 min, vertical scale bar: 10 μM. **(F)** Left: Schematic showing the domain organization of the Ufd1 protein. Asterisks indicate the location of the mutations present in *ufd1-1* and *ufd1-5*, respectively. Right: Tetrads from crosses between *cdc48-353* and *ufd1* mutants, *ufd1-1* (left) and *ufd1-5* (right). Colored circles indicate the strain genotype, dashed colored circles indicate spore lethality.

Since the phenotype caused by Cdc48G338D is rescued by impairing the D1 ATPase activity as well as by mutating or deleting cofactors that cooperate with Cdc48, we conclude that *cdc48-353* is a gain-of-function allele whose toxic effects are mitigated by making Cdc48 less active. These findings further conflict with the proposed model, which assumes *cdc48-353* to be a loss-of-function allele (Fig. 1A).

### The rescue of the *cdc48-353* mutant by deletion of the Ufd2 cofactor is independent of securin degradation during mitosis

Deletion of the Cdc48 cofactor *ufd2^+^* strongly suppresses the *cdc48-353* mutant growth phenotype (Fig. 5A) (Xu et al., 2018). Accordingly, we found that separase, securin and Cdc13 levels were restored in the *cdc48-353 ufd2Δ* double mutant (Fig. 5A-C; S5B). Ufd2 is an E4 ubiquitin ligase that in budding yeast has been shown to add ubiquitin moieties to preformed ubiquitin conjugates to aid proteasomal degradation (Richly et al., 2005; Rumpf and Jentsch, 2006). The suppression observed in the *cdc48-353 ufd2Δ* double mutant has been proposed to arise from inefficient securin degradation rescuing separase from undergoing co-degradation (Xu et al., 2018). If this was true, we would expect inefficient securin degradation in the *ufd2Δ* mutant. However, despite slightly higher securin and separase levels in the *ufd2Δ* mutant, we found no obvious change in securin degradation kinetics during mitosis (Fig. 5D,E; S5B,C). These results indicate that the rescue of separase levels seen in the *cdc48-353 ufd2Δ* double mutant is unlikely to be a consequence of impaired securin degradation during mitosis.

A *cdc48-353* suppressor screen identified a frameshift mutation in *ufd1^+^* (Marinova et al., 2015), another Cdc48 cofactor. Ufd1 in complex with Npl4 binds ubiquitinated substrates and helps Cdc48-mediated unfolding and processing (Meyer et al., 2002; Rape et al., 2001; Rumpf and Jentsch, 2006; Twomey et al., 2019). We tested the genetic interaction of two *ufd1* mutants (*ufd1-1* and *ufd1-5*) with *cdc48-353*. The first of these (*ufd1-1*, P76S) is expected to impair ubiquitin binding, the second (*ufd1-5*, L218S) is expected to impair Cdc48 binding (Burr et al., 2017; Hänzelmann and Schindelin, 2016; Nie et al., 2012; Twomey et al., 2019). The *ufd1-1* mutant was synthetic lethal with *cdc48-353* whereas the *ufd1-5* mutant partially rescued *cdc48-353* (Fig. 5A, B,F). This indicates that interaction of Ufd1 with Cdc48 is important for the toxic effect of *cdc48-353*. If securin degradation was indeed facilitated by Ufd1/Npl4-Cdc48, we would expect securin overexpression to be toxic in *ufd1* mutants. However, securin overexpression was tolerated by both the *ufd1-1* and *ufd1-5* mutants (Fig. S5F).

In summary, these results show that the detrimental effects of *cdc48-353* can be alleviated by removing Cdc48 cofactors or impairing their interaction with Cdc48. This supports the idea that *cdc48-353* is a gain-of-function allele. The fact that deletion of *ufd2^+^* strongly rescues the *cdc48-353* mutant phenotype, but does not impair securin degradation, further supports our conclusion that low separase levels are independent of mitotic securin degradation and not a consequence of co-degradation of separase with securin in the *cdc48-353* mutant.

### TORC1 mutation rescues *cdc48-353* growth and mitotic defects without restoring separase levels

In addition to cofactor deletions, growth on minimal medium suppresses the temperature sensitivity of the *cdc48-353* mutant suggesting a link to nutrient availability (Ikai and Yanagida, 2006). The conserved TOR (target of rapamycin) pathway controls cell growth in response to nutrients (González and Hall, 2017). Mutations in TOR complex 1 (TORC1) have been previously shown to rescue the phenotype of separase mutants (Ikai et al., 2011). We therefore asked whether a mutant of *tor2*, which is part of TORC1, would also rescue the effects of *cdc48-353*. The Tor2S1837E mutation (*tor2SE;* (Laor et al., 2014)) indeed rescued *cdc48-353* growth defects as well as mitotic phenotypes, such as chromosome bridges and the ‘cut’ phenotype (Fig. 6A,B). Strikingly, though, separase and securin levels remained unaltered (Fig. 6C; S6A). Consistently, we found that sister chromatid separation remained delayed in *tor2SE cdc48-353* cells (Fig. S6B,C). Although surprising, this fits with the prior observation that growth of the separase mutant *cut1-206* is rescued by the TOR inhibitor rapamycin although separase levels are not restored (Ikai et al., 2011). A phenotype of shorter mitotic spindles that we find in the *cdc48-353* mutant, but not in the separase *cut1-206* mutant, was also not efficiently rescued by *tor2SE* (Fig. S6D). Hence, reduced TOR signalling specifically rescues the mitotic cut phenotype without restoring separase levels.

**Figure 6.**
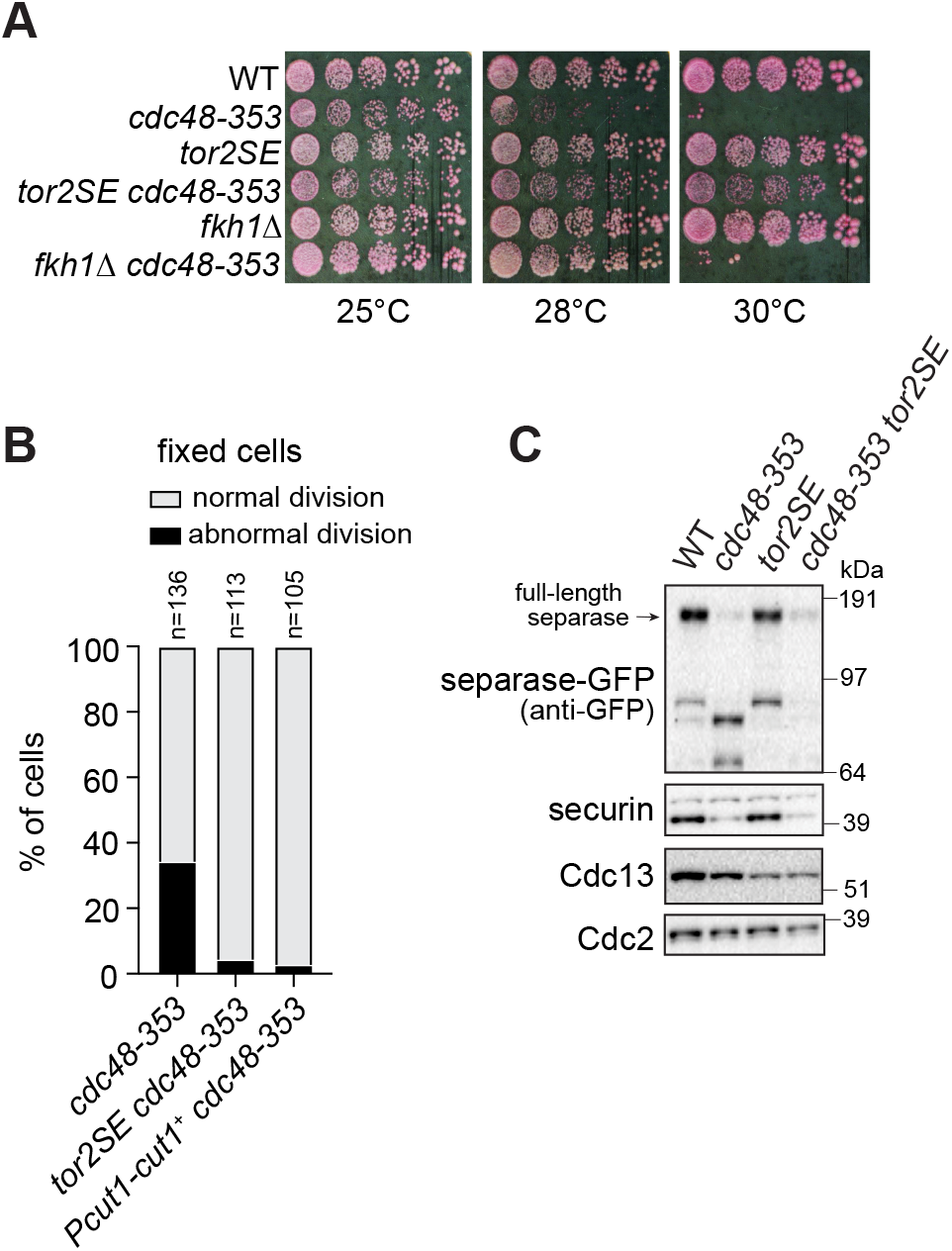
Mutation of *tor2* rescues both growth and mitotic defects in the *cdc48-353* mutant without rescuing separase levels. **(A)** Growth of wild type (WT) and the indicated single and double mutant strains at different temperatures on rich medium with Phloxine B, which stains dead cells. **(B)** Phenotypic analysis of the indicated strains by DNA staining of fixed cells. Abnormal divsion was scored by counting cells with cut phenotype, unequally divided nucleus or chromosome bridges. The *cdc48-353* data are the same as in Figure 7D. **(C)** Immunoblot showing the levels of the indicated proteins in WT, *cdc48-353, tor2SE* and *tor2SE cdc48-353* double mutant.

Nuclear division in *S. pombe* cells requires nuclear membrane expansion, and mutations or drugs that impair lipid synthesis also cause a cut phenotype (Saitoh et al., 1996; Takemoto et al., 2016; Yam et al., 2011). Such mutants are also rescued by minimal medium—more specifically by the ammonium present in standard minimal medium (Zach et al., 2018). Altogether, this suggests that *cdc48-353* and separase mutants may be defective in lipid metabolism, which impairs mitosis and can be rescued by ammonium or reduced TORC1 signalling.

In summary, TORC1 mutation rescues some aspects of the *cdc48-353* mitotic phenotype, but without rescuing separase levels. This establishes that the lowering of separase levels is not the only pertinent defect in *cdc48-353* cells and that effects other than low separase levels contribute to the growth defect and failed mitosis.

### Low separase levels in the *cdc48-353* mutant do not depend on any major degradation pathway

If low separase levels in the *cdc48-353* mutant were a consequence of mitotic co-degradation with securin (Fig. 1A), a rescue of separase levels would be expected when impairing degradation. However, combining *cdc48-353* with a proteasome mutant (*mts3-1*) (Ikai and Yanagida, 2006) or chemical inhibition of proteasome activity by Velcade (bortezomib) failed to rescue separase levels in the *cdc48-353* mutant (Fig. 1B, 7B). In contrast, the levels of securin and Cdc13 increased, indicating that different mechanisms are behind the low levels of these three proteins in the *cdc48-353* mutant. The results suggest that the low separase levels are not a direct consequence of proteasomal degradation.

**Figure 7.**
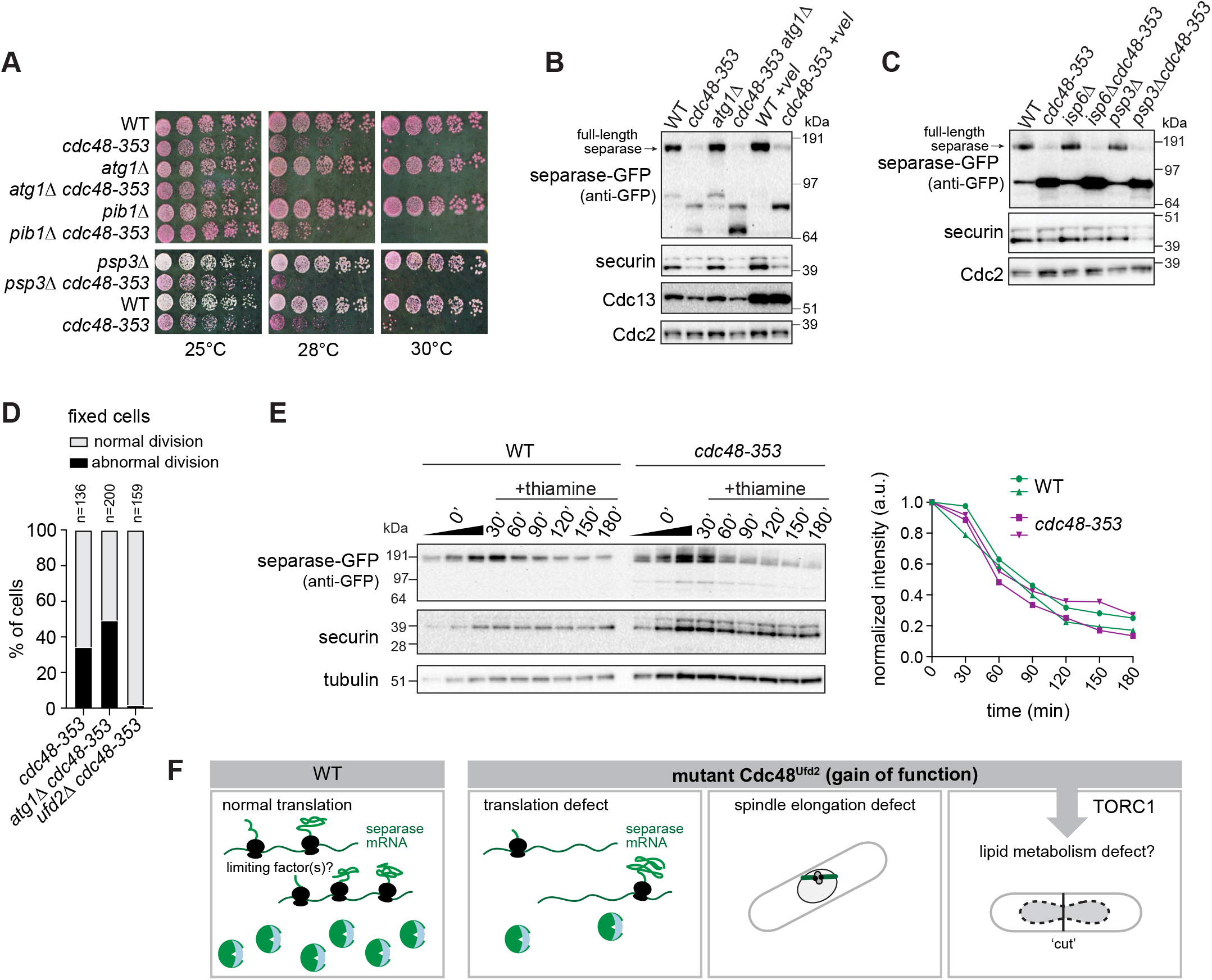
Separase stability is unaltered in the *cdc48-353* mutant and inhibition of major degradation pathways fails to rescue separase levels. **(A)** Growth of wild type (WT) and the indicated single and double mutant strains at different temperatures on rich medium with Phloxine B, which stains dead cells. **(B)** Immunoblot showing levels of the indicated proteins in WT and *cdc48-353* mutant strains with or without inhibition of autophagy by *atg1* deletion or inhibition of the proteasome by velcade (+vel) treatment. Higher levels of Cdc13 upon velcade treatment suggest efficacy of proteasomal inhibition. **(C)** Immunoblot showing levels of the indicated proteins in WT and *cdc48-353* mutant strains with or without deletion of the vacuolar proteases Isp6 and Psp3. **(D)** Phenotypic analysis of the indicated strains by DNA staining of fixed cells. Abnormal divsion was scored by counting cells with cut phenotype, unequally divided nucleus or chromosome bridges. The *cdc48-353* data are the same as in Figure 6B. **(E)** *Nmt81* promoter-driven expression of separase-GFP in WT and *cdc48-353* mutant cells at the indicated time points after thiamine addition to shut off transcription. Because separase levels are low in the *cdc48-353* mutant, approx. 4 times more extract than for WT cells was loaded for better comparability. Left: Immunoblot, with 1:1 dilution series loaded for the 0’ time point before addition of thiamine. Right: Quantification of two independent experiments. **(F)** Model proposing that the low separase levels in the *cdc48-353* mutant are a result of poor separase translation, and that the *cdc48-353* mutation impairs mitosis in multiple ways.

This prompted us to test alternative degradation pathways. Autophagy was a particularly promising candidate since Cdc48 has been shown to promote autophagy (Dargemont and Ossareh-Nazari, 2012; Franz et al., 2014). However, deletion of the core autophagy kinase *atg1^+^* neither rescued growth of the *cdc48-353* mutant, nor separase levels or abnormal mitosis (Fig. 7A,B,D). Similarly, deletion of the vacuolar ubiquitin ligase *pib1^+^* or the vacuolar serine proteases *isp6^+^* and *psp3^+^* did not rescue growth or separase levels (Fig. 7A,C). Hence, we cannot positively confirm any of the major cellular degradation pathways to be involved in lowering separase levels in the *cdc48-353* mutant.

We did observe two prominent faster migrating bands in the GFP immunoblot of *cdc48-353* cells expressing separase-GFP (Fig. 5B, 6C, 7B). These bands were affected by interfering with the different degradation pathways, even when the full-length band was not (Fig. 7B), and they disappeared upon combination of *cdc48-353* with the *tor2SE* mutant (Fig. 6C). However, these bands seemed absent in cells overexpressing the mutant Cdc48G338D (Fig. 4B) and did not appear at corresponding sizes in *cdc48-353* mutant cells expressing myc-tagged separase (Fig. S2D). Hence, the identity and possible role of these shorter fragments remains unclear.

Overall, these results raise doubts whether separase levels in the *cdc48-353* mutant are lowered by degradation at all, or if synthesis may be impaired.

### Low separase levels may result from poor separase translation in the *cdc48-353* mutant

In order to more generally address whether separase is becoming destabilized in the *cdc48-353* mutant, we assessed separase levels after promoter shut-off, using a second copy of separase integrated at the *leu1* locus. Separase mRNA has a short half-life (3.6 min; (Eser et al., 2016)), so that a transcriptional shut-off is expected to rapidly stop protein production. The timing of separase decline was similar for wild-type and mutant cells (Fig. 7E). This provides further evidence that separase is not destabilized in the *cdc48-353* mutant and instead suggests that separase is less efficiently produced in the *cdc48-353* mutant. Separase mRNA levels have been shown to be unaffected in the *cdc48-353* mutant (Ikai and Yanagida, 2006), and we therefore speculate that translation is impaired. Based on recent genome-wide ribosome profiling in *S. pombe*, separase has a very low translation efficiency (Fig. S7A), which makes it plausible that its translation requires specific factors that become limiting in *cdc48-353* cells (Fig. 7F).

Overexpression of Int6 and Moe1, components of the translation initiation complex eIF3, have been reported to rescue *cdc48-353* growth; and *cdc48-353* defects are further aggravated by *int6^+^* or *moe1^+^ deletion* (Otero et al., 2010). To understand the mode of this rescue we overexpressed Int6 in wild-type and *cdc48-353* mutant strains. However, separase levels remained unchanged upon Int6 overexpression in the *cdc48-353* mutant (Fig. S7B) and we could not confirm the rescue of growth defects upon Int6 overexpression from its endogenous promoter or from a strong *nmt1* promoter (Fig.S7C,D).

Altogether, these results argue for a separase translation defect in cdc48-353 mutant cells, and therefore that separase has specific requirements for its translation that are not generally shared by all cellular proteins (Fig. 7F). What factors aid separase translation remains enigmatic.

## Discussion

In contrast to the long-standing hypothesis that separase undergoes mitotic co-degradation with securin in the *S. pombe cdc48-353* mutant, which would have implicated Cdc48 in the regulation of anaphase (Fig. 1A), we show here that low separase levels in this *cdc48* mutant occur independent of securin degradation in anaphase and that Cdc48 does not play a major role in securin degradation and separase activation during mitosis (Fig. 3A). Prior experiments in *Xenopus laevis* extracts reached the same conclusion (Heubes, 2007). The unfoldase activity of Cdc48 is thought to be critical for proteasomal degradation when substrates are well-folded and lack a flexible region (Beskow et al., 2009; Olszewski et al., 2019). Securin, in contrast, is a largely unstructured protein whose ubiquitinated N-terminus is natively unfolded (Cox et al., 2002; Csizmok et al., 2008; Sánchez-Puig et al., 2005). It is therefore plausible that securin is efficiently degraded without the help of Cdc48. Separase becomes activated not only during mitosis, but also locally in interphase upon DNA damage (Hellmuth et al., 2018; McAleenan et al., 2013; Nagao et al., 2004). Cdc48 is known to be involved in DNA repair (Franz et al., 2016; Torrecilla et al., 2017). Whether Cdc48 plays a role in temporarily and locally removing securin from separase in this situation remains an open question.

Our results show that the *cdc48-353* phenotype is more multi-faceted than previously appreciated (Fig. 7F). Not all aspects of the mitotic phenotype can be attributed solely to the reduction in separase levels—for example, the short prometaphase spindles seen in *cdc48-353* cells are not observed in a separase mutant (Fig. S6). Whereas deletion of the Cdc48 cofactor *ufd2^+^* rescued cell growth, separase levels and mitotic phenotype, mutation of *tor2* rescued the mitotic cut phenotype but did not restore separase levels. This suggests that low separase levels are not the sole reason for the cut phenotype of *cdc48-353* cells. Alternatively, the *tor2SE* mutation may alleviate the need for separase for faithful completion of mitosis. We therefore tried if separase can be deleted in the *cdc48-353 tor2SE* double mutant but failed to obtain such strains (data not shown). This suggests that separase remains necessary for successful mitosis, and that the low levels of separase in the *cdc48-353* mutant are not the only reason why these cells fail to divide properly.

The current data suggest that *cdc48-353* is a gain-of-function mutation, and we propose that the mutant Cdc48, along with its cofactors Ufd1 and Ufd2, over-processes one or more natural substrates, thereby impairing their function. The involvement of the Ufd1 and Ufd2 cofactors suggests that a relevant substrate is likely ubiquitinated. A comparative proteome analysis or comparative Cdc48 immunoprecipitations between *cdc48-353* and wild-type cells may identify the relevant substrates. As of now, there is no evidence that Cdc48 acts directly on securin and separase, and we propose that the effect of lowering separase levels in the *cdc48-353* mutant is indirect. In contrast to the previously prevailing hypothesis that separase has become unstable in the *cdc48-353* mutant, our data suggest that separase is inefficiently produced.

Separase requires securin for its stability and full activity (Hornig et al., 2002; Jallepalli et al., 2001; Rosen et al., 2019) and securin starts to associate with separase during separase translation (Hellmuth et al., 2015). Hence a failure of establishing this interaction may lead to reduced separase translation. However, we did not find a defect in securin-separase interaction in the *cdc48-353* mutant (Fig. S3A). It remains possible, though, that co-translational association is inefficient and leads to a defect in separase translation, but that the few successfully formed securin-separase complexes are fully translated and only those are evaluated in our assay. Given the stabilizing effect of securin, we also considered whether the low separase levels could be a consequence of a failure to sustain proper concentrations of securin in the *cdc48-353* mutant. However, this seems unlikely given that expression of additional securin is toxic and does not rescue separase levels (Fig. 2) (Ikai and Yanagida, 2006; Yuasa et al., 2004).

Alternatively, factors other than securin association may be missing for separase translation in the *cdc48-353* mutant. In addition to the canonical translation machinery, non-canonical transacting proteins as well as cellular tRNA composition affect the translation of specific subsets of mRNAs (Baltz et al., 2012; Harvey et al., 2018; Rak et al., 2018). The previously established genetic interaction between *cdc48* and components of the eIF3 translation initiation complex (Otero et al., 2010) and findings that these eIF3 components promote proper chromosome segregation (Yen and Chang, 2000) were exciting, but we have so far been unable to verify this connection (Fig. S7). Cdc48 has also been shown to promote the disassembly and subsequent degradation of defective or stalled RNA polymerase III complexes (Wang et al., 2018). Hence, it is possible that *cdc48-353* shapes the tRNA pool with consequences on the translation of separase and likely other proteins.

Although it remains unknown how mutant Cdc48 impairs separase, our study suggests that separase production has specific requirements that can be impaired by dysfunctional Cdc48 leading to catastrophic cellular outcomes. It will be interesting to address what these requirements are and what purpose they serve in wild-type cells. This may also shed light on the causes of the separase overexpression that is often seen in cancer cells (Meyer et al., 2009).

## Materials and Methods

### *S. pombe* strains

Strains used in this study are listed in Table S1. Tagging or deletion of genes at the endogenous locus was performed by PCR-based gene targeting (Bähler et al., 1998). For the mutagenesis screen of securin, the *cut2^+^* coding sequence was mutagenized by PCR using Taq polymerase and conditions that further compromise fidelity of the polymerase (Wilson and Keefe, 2001). The mutant library was introduced into the pREP1-Pnmt1-GFP vector backbone by Gibson assembly (New England Biolabs). In this vector, *cut2* was fused to GFP at its C-terminus and expressed under the *nmt1* promoter (*Pnmt1*). The validation of the *cut2* mutagenesis screen was performed by introducing the indicated *cut2* variants into a pREP81 vector where they are expressed under the weaker *nmt81* promoter (a mutated version of the *nmt1* promoter). For strains expressing a second copy of separase, a pDUAL vector (Matsuyama et al., 2004) containing *Pcut1-cut1^+^-GFP* or *Pcut1-cut1^+^-(myc)_13_* was linearized with NotI and integrated at the *leu1* locus. 901 bp of *cut1* region upstream of its start codon were used. For strains expressing *cut1^+^-GFP* under the regulatable *nmt81* promoter, *pDUAL-Pnmt81-cut1^+^-GFP* was linearized with NotI and integrated at the *leu1* locus. For plasmids expressing *cdc48^+^* and mutant versions, a (his)6 tag was introduced at the N-terminus of *cdc48* by PCR and the entire fragment was introduced into NdeI/BamHI digested pREP1-Pnmt1 vector. To generate the *Pint6-int6^+^* expression construct, full-length *int6^+^* including 1011 bp upstream of the *int6^+^* start codon was inserted into the pREP1 vector replacing the *nmt1* promoter. For *Pnmt1* driven overexpression of *int6^+^*, full-length *int6^+^* was fused to a dual flag tag at the C-terminus in a pDUAL vector which was linearized with NotI and integrated at the *leu1* locus. For the Cdc48 anchor away strains, Cdc48 was C-terminally tagged with either FRB-GFP or only FRB (for strains expressing Cut1-GFP). Ribosomal protein Rpl13 tagged with 2 tandem copies of FKBP12 at the C-terminus was integrated at the *leu1* locus and expressed from the *Pnmt1* promoter.

### Cell culture

For all immunoblots and imaging assays, cells were cultured in Edinburgh minimal medium (EMM) with supplements for auxotrophic strains as needed (Moreno et al., 1991). To suppress expression from *nmt1* or *nmt81* promoters, 16 *μ*M thiamine was used. For growth assays, cultures were grown in EMM media containing 16 *μ*M thiamine, followed by thiamine washout before spotting on EMM plates with or without thiamine. Growth assays on rich media were on yeast extract with adenine (YEA) including 2 *μ*g/mL phloxine B. To inhibit proteasome activity, bortezomib (Velcade, B-1408, LC Laboratories) was added at a final concentration of 1 mM to cultures in exponential growth phase. Cultures were incubated at 30°C for 45 minutes in the presence of the drug. A small volume of the culture was fixed with methanol to check for proteasomal inhibition by scoring for cells arrested at metaphase. Rapamycin (R-5000, LC Laboratories) treatment was with 2.5 *μ*g/mL for 45 minutes prior to the start of imaging.

### *Pnmt81* promoter shut down of separase expression

WT and *cdc48-353* mutant stains expressing *Pnmt81-cut1^+^-GFP* from the *leu1* locus were grown in EMM medium lacking thiamine for 15 hours to induce Cut1-GFP expression; 1 x 10^8^ cells were harvested from both WT and mutant cultures, 16 *μ*M thiamine was added to both cultures and 1 x 10^8^ cells were harvested at every 30-minute interval for 3 hours. Harvested cells were washed with 20 % trichloroacetic acid, spun down at 1,150 rcf and snap-frozen in liquid nitrogen. Protein samples were prepared using the denatured extract preparation described below.

### Live-cell imaging

For live-cell imaging, cells were mounted in lectin-coated (40 *μ*g/ml; Sigma L1395) culture dishes (8-well; Ibidi) at a density of 1.5 x 10^6^ cells/ml and pre-incubated on the microscope stage at 30°C for 45 minutes. For temperature-sensitive strains, the culture was grown at 25°C and cells were shifted to 30°C when pre-incubating on the microscope stage for 30 minutes prior to imaging at 30°C for 90 minutes. Images were acquired using a 60x/1.4 Apo oil objective (Olympus) on a DeltaVision Elite system (Applied Precision/GE Healthcare) equipped with a PCO edge sCMOS camera and an environmental chamber for temperature control.

### Image acquisition and analysis

Images were acquired using the optical axis integration (OAI) mode available with the SoftWorx software using a range of 3.6 *μ*M. Images were acquired every 15 seconds for 90 minutes to follow the levels of GFP-tagged separase, securin and Cdc13 in cells undergoing mitosis. SCS in these cells was scored by their dh1L<<tetO/TetR-tdTomato signal (Sakuno et al., 2009). Fiji/ImageJ (Schindelin et al., 2012) was used to quantify nuclear and cellular concentrations of the protein analyzed. Nuclear and cellular region of interest (ROI) at each time point were defined using TetR-tdTomato- and brightfield-based thresholding, respectively. The mean background outside of cells was subtracted. The degradation kinetics were determined in Matlab (MathWorks). A cubic spline was fit to the degradation curve. The onset of degradation was defined as the point when the degradation rate first reached 10 % of the maximal degradation rate; the end of degradation was defined as the point when the curve first became flat. Data was normalized to the intensities at these two points. For the normalized degradation rate, the region of the curve between 30 and 70 % of degradation was used to determine the slope.

### Cell extracts and immunoblotting

Immunoblotting used denatured extracts or native extracts. For denatured extracts, protein samples were prepared using trichloroacetic acid-based extraction: 1 x 10^8^ cells were resuspended in 500 *μ*L water. To this suspension, 75 *μ*L of 1.85 M sodium hydroxide/1 M β-mercaptoethanol was added and the mixture incubated on ice for 15 minutes. 75 *μ*L of 55 % trichloroacetic acid were added and incubation continued on ice for another 15 minutes. The samples were spun down at 16,000 rcf for 15 minutes and supernatant was completely removed from the pellet. The pellets were resuspended in 180 *μ*L of sample buffer (500 *μ*L of HU buffer [8 M urea, 5 % SDS (w/v), 200 mM Tris-HCl pH 6.8, 20 % glycerol (v/v), 1 mM EDTA, 0.1 % (w/v) bromophenol blue], 400 *μ*L of 2 M Tris-HCl pH 6.8, 100 *μ*L of 1 M DTT). 150 *μ*L of acid-washed beads (G8772, Sigma) were added to the resuspended pellet and tubes agitated in a ball mill (RETSCH MM400) for 2 minutes at 30 Hz. Tubes were pierced at the bottom and eluates collected into fresh tubes by centrifugation at 2,400 rcf for 1 minute. The extracts were heated at 75°C for 5 minutes. Lysate corresponding to 5 x 10^6^ cells was loaded per lane. Proteins were separated by SDS-PAGE (4-12 % Bis-Tris NuPAGE gel, Invitrogen) and the resolved proteins were transferred onto PVDF membranes (Immobilon-P, Millipore) using a semi-dry blotting assembly (Amersham Biosciences) with Trisglycine buffer containing 10 % methanol and 0.01% SDS.

For native extracts, cell pellets were resuspended in 100-250 *μ*L of sterile water and frozen as droplets in liquid nitrogen. Cells were disrupted under cryogenic conditions using a ball mill (RETSCH MM400) for 15 seconds at 30 Hz. A small sample of the protein powder was resuspended in lysis buffer (20 mM Tris-HCl (pH 7.5), 150 mM NaCl, 5 % glycerol and 0.1 % NP-40) to determine the protein concentration by BCA assay (Thermo Scientific). For experiments, frozen powder was resuspended to a final concentration of 10 *μ*g/*μ*L in lysis buffer supplemented with Halt protease and phosphatase inhibitor cocktail (Thermo Scientific). Extracts were spun down at 16,000 rcf for 10 minutes at 4°C and the supernatant was collected. For immunoblotting, the supernatant was mixed with an equal volume of sample buffer (900 *μ*L HU buffer + 100 *μ*L of 1 M DTT) and heated to 75°C for 5 minutes. 30 *μ*g of protein was loaded into one lane.

Primary antibodies were goat anti-p97 (Abcam, ab206320), mouse anti-myc (Sigma, M4439), mouse anti-Cdc13 (cyclin B, Novus, NB200-576), rabbit anti-Cdc2 (CDK1,Santa Cruz, SC-53), mouse anti-tubulin (Sigma, T5168), rabbit anti-Cut2 (Kamenz and Hauf, 2014), and mouse anti-GFP (Roche, 11814460001). Secondary antibodies were anti-mouse or anti-rabbit HRP conjugates (Dianova, 115-035-003, 111-035-003) and anti-goat HRP conjugate (Abcam, ab6741). Blots were developed using an enhanced chemiluminescent substrate (SuperSignal West Pico Plus, ThermoFisher).

### Immunoprecipitation

Immunoprecipitations of securin and separase were performed using native extracts prepared as above. Mouse anti-GFP monoclonal antibody (Roche, 11814460001) was covalently coupled to magnetic beads at 12 *μ*g antibody per 100 *μ*L Dynabeads Protein A (ThermoFisher). Around 1500 *μ*g of cell lysate and 10 *μ*g of antibody was used per sample. Immunoprecipitates were eluted using 7 *μ*l of 100 mM citric acid pH 2; pH of the sample was adjusted with 1.4 *μ*l of 1 M Tris pH 9.2, and the elution boiled with 8.6 *μ*l of HU buffer (8 M urea, 5 % SDS, 200 mM Tris HCl pH 6.8, 1 mM EDTA, 20 % glycerol and 0.01 % bromophenol blue).

### Calculating translation efficiency

Translation efficiency was calculated as the ratio between previously reported raw counts of ribosome-protected RNA fragments and RNA-seq of wild-type untreated cells from different experiments by Rubio et al., 2020 (Rubio et al., 2021). Only non-mitochondrial, non-dubious protein-coding genes were included, which resulted in a list of 5,057 genes. From this, an additional 30 genes were excluded because they had no RNA-seq raw counts or data was only available from four or fewer experiments out of a total of ten experiments.

## Acknowledgments

We thank Tatiana Boluarte for help with yeast strain construction, Michael Boddy, Peter Espenshade, Yoshinori Watanabe, and Mitsuhiro Yanagida for providing yeast strains, Susan Forsburg for providing plasmids, as well as Andrea Ciliberto, Julia Kamenz, and all members of the Hauf Lab for critical reading of the manuscript. This work was supported by the NIH/National Institute of General Medical Sciences under award number R35GM119723.

## Author contributions

Conceptualization: DV, SH; Investigation and Visualization: DV, JM; Funding acquisition: SH; Writing: DV, SH.

## Competing Interest

None declared.

## Notes

### Competing Interest Statement

The authors have declared no competing interest.

