## Supplemental Figures and Table for "Cdc48 influence on separase levels is independent of mitosis and suggests translational sensitivity of separase"

### Figure S1
Vijayakumari et al.

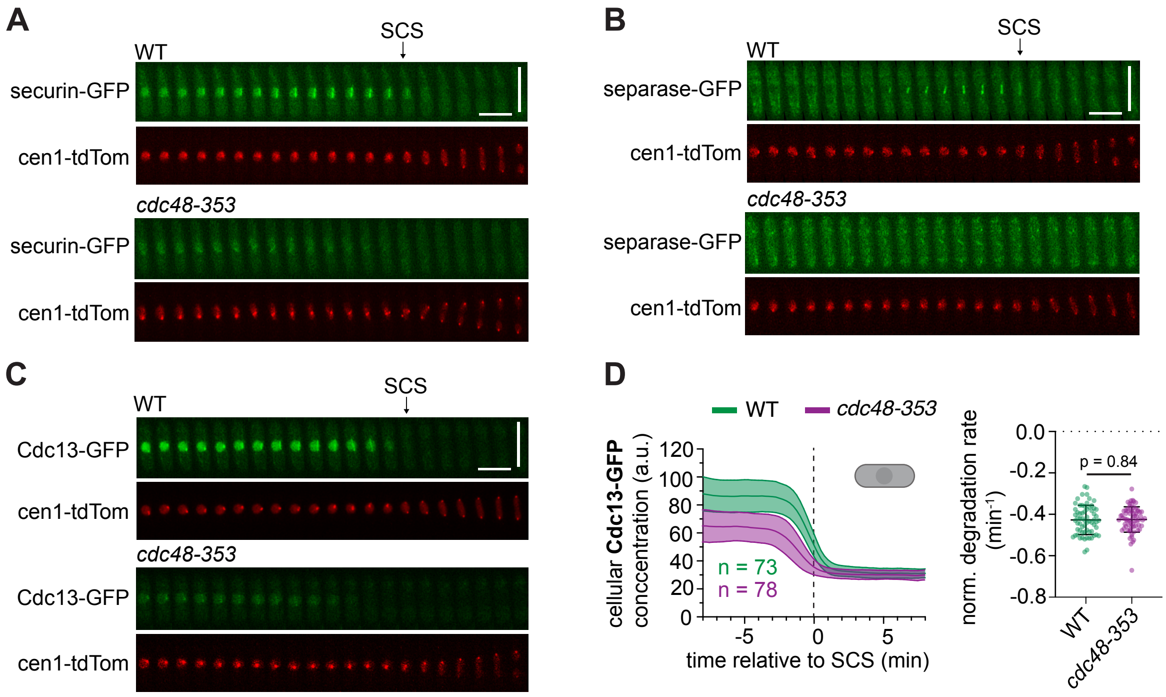

**Figure S1. Reduced levels of separase, securin and Cdc13 (cyclin B) in the *cdc48-353* mutant during mitosis. (A-C)** Kymographs showing securin-GFP (A), separase-GFP (B), and Cdc13-GFP (C) in the wild type (WT) and *cdc48-353* mutant as cells progress through mitosis; tdTomato at the centromere of chromosome 1 (cen1-tdTom) serves as marker for sister chromatid separation (SCS). Horizontal scale bar: 2 min, vertical scale bar: 10 µm. **(D)** Left: Cellular concentration of Cdc13(cyclin B)-GFP in the wild type (WT) and *cdc48-353* mutant during mitosis. The data are aligned to the time point of sister chromatid separation (SCS), indicated by the vertical dashed line. Right: The normalized degradation rate of Cdc13 during mitosis. Bars are mean and standard deviation. Statistical significance was tested by unpaired t-test.

**Figure S2**
**Vijayakumari et al.**

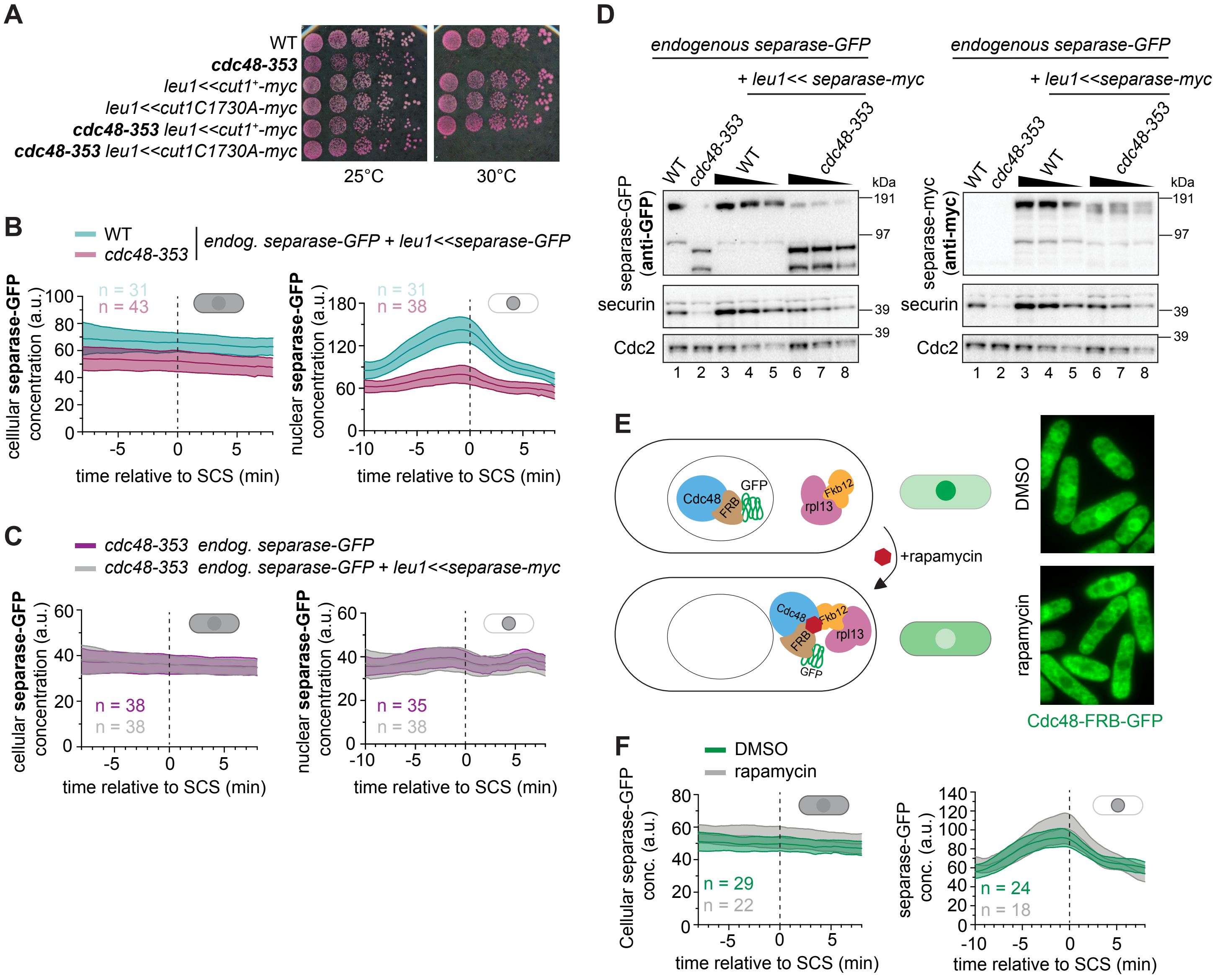

**Figure S2. Separase expressed from the *leu1* locus is affected by the *cdc48-353* mutant. (A)** Growth assay of wild-type (WT) and *cdc48-353* mutant strains expressing a second copy of wild-type separase or protease dead (C1730A) separase integrated at the *leu1* locus. WT and *cdc48-353* mutant without the second copy of separase (first two rows) serve as control. Cells were plated on rich medium with Phloxine B, which stains dead cells. **(B)** The cellular (left) and nuclear (right) concentration of separase-GFP in WT or *cdc48-353* mutant cells expressing two copies of separase-GFP. The data for *cdc48-353* are the same as in Figure 3. **(C)** The cellular (left) and nuclear (right) concentration of separase-GFP in the *cdc48-353* mutant either without or with an additional copy of myc-tagged separase expressed from the *leu1* locus. The data without extra separase are the same as in Figure 3. **(D)** Immunoblots showing the levels of separase-GFP expressed from the endogenous locus (anti-GFP) and separase-myc expressed from the *leu1* locus (anti-myc). Lanes 1 and 2 contain extracts of strains only expressing separase-GFP from the endogenous locus. Lanes 3 - 8 contain extracts of strains expressing both separase-GFP from the endogenous locus and separase-myc from the *leu1* locus. Securin and Cdc2 levels in these strains are also shown. Serial 1:1 dilutions of some extracts are indicated by the wedge. The expression of separase-GFP at the endogenous locus is unaffected by the second copy of separase-myc (left side). The level of separase-myc expressed from the *leu1* locus is also lowered by the *cdc48-353* mutation (right side). **(E)** Left: Schematic showing genotype of the strain used to conditionally anchor away Cdc48 from the nucleus in the presence of rapamycin. Right: Representative images showing the exclusion of Cdc48-FRB-GFP from the nucleus upon treatment with rapamycin (2.5 µg/ml for 45 minutes). **(F)** The cellular (left) and nuclear (right) concentration of separase-GFP during mitosis in a strain where Cdc48 is anchored away from the nucleus after treatment with rapamycin (grey). The DMSO-treated control is shown in green. Note that Cdc48 is not tagged in this experiment, so it remains uncertain whether nuclear Cdc48 levels are as efficiently reduced as in (E).

**Figure S3**
**Vijayakumari et al.**

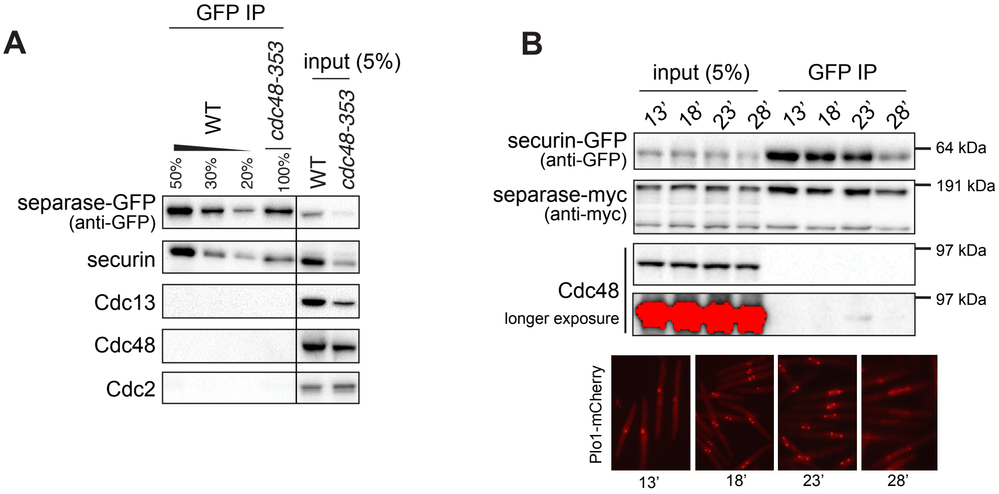

**Figure S3. No strong physical interaction between separase and Cdc48 in asynchronous cultures or between securin and Cdc48 during mitosis. (A)** Separase-GFP immunoprecipitation (IP) from asynchronous cultures of wild type (WT) and *cdc48-353* mutant. **(B)** Securin-GFP IP from *cdc25-22* cells at different time points during mitosis after release from G2/M arrest. Plo1-mCherry was used to asess progression through mitosis and representative images corresponding to each time point are shown. Red pixels in the immunoblot indicate saturation of the camera.

### Figure S4
### Vijayakumari et al.

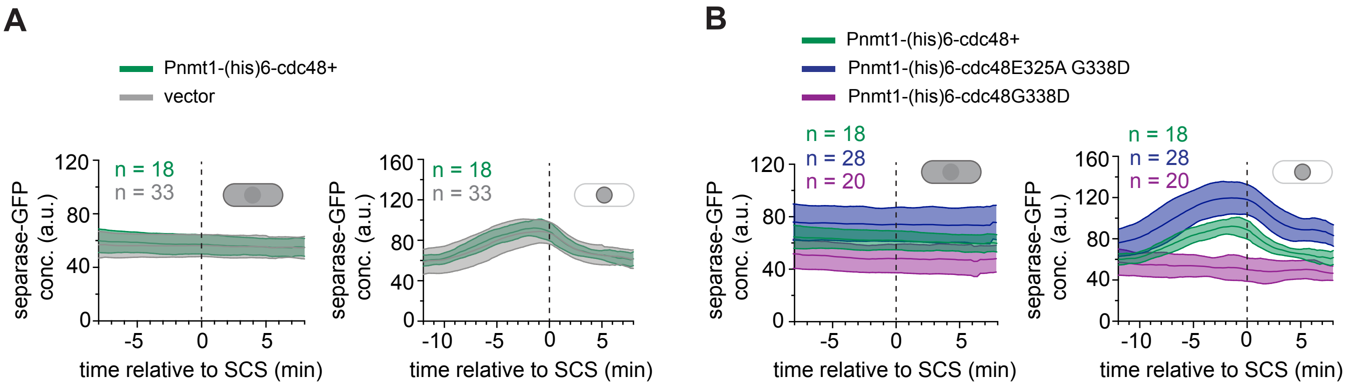

**Figure S4. Wild-type cells overexpressing mutant Cdc48G338D have low separase levels. (A)** Comparison of the cellular (left) and nuclear (right) concentration of separase-GFP during mitosis in wild-type cells overexpressing Cdc48 (green) or containing an empty vector as control (grey). Mean (central line) and standard deviation (area) are shown (n = number of cells). **(B)** Comparison of the cellular (left) and nuclear (right) concentration of separase-GFP during mitosis in wild-type cells overexpressing *cdc48*$^{+}$, *cdc48G338D* or *cdc48E325A G338D*.

**Figure S5**
**Vijayakumari et al.**

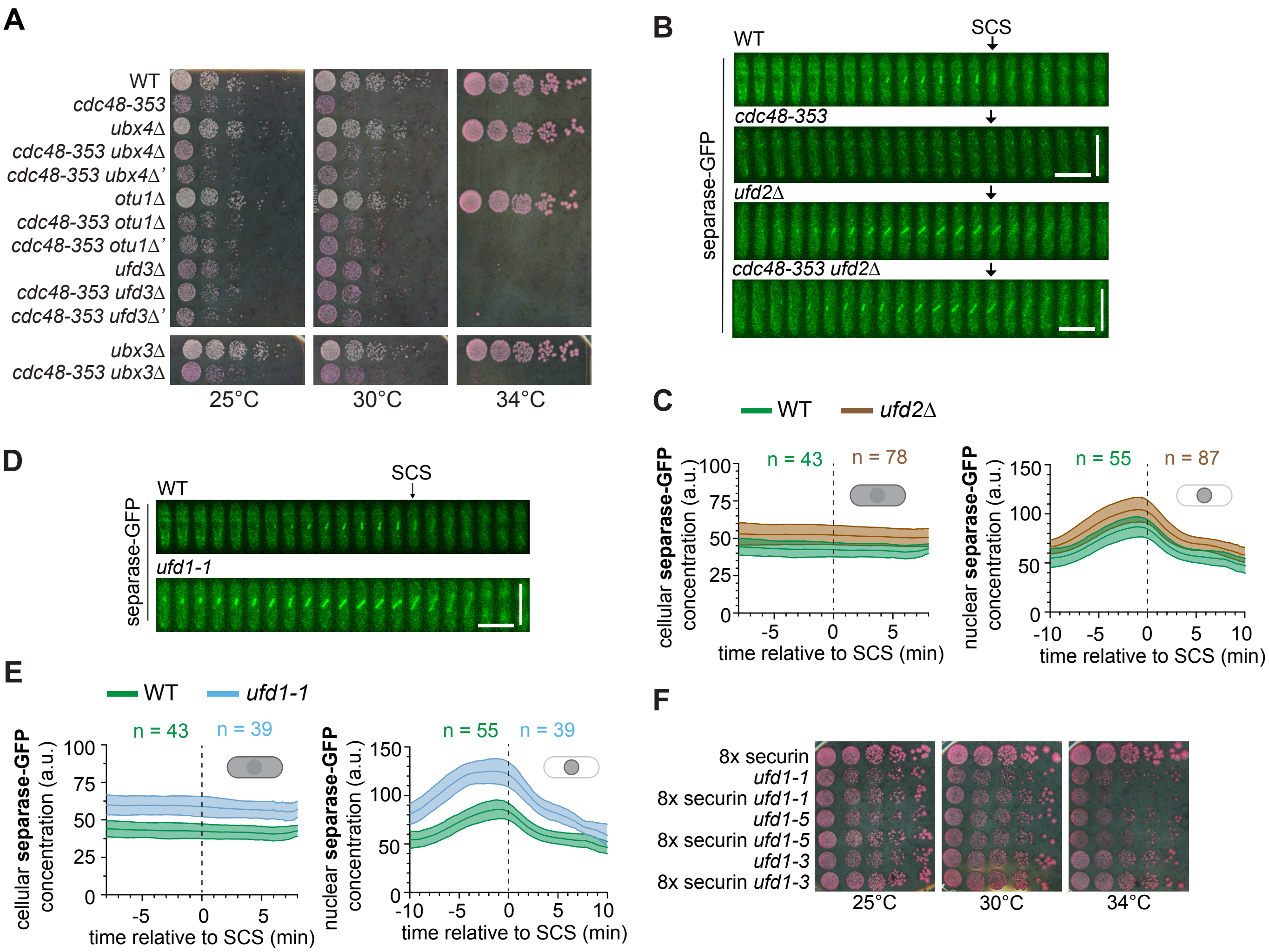

**Figure S5. Separase levels are rescued in the *cdc48-353 ufd2*Δ double mutant and are increased in *ufd1-1*. (A)** Growth assay of wild type (WT), *cdc48-353* single mutant and double mutant combinations with different Cdc48 co-factor deletions at the indicated temperatures. Cells were plated on rich medium with Phloxine B, which stains dead cells. **(B)** Representative kymographs showing separase-GFP in wild-type (WT), *cdc48-353*, *ufd2*Δ and *cdc48-353 ufd2*Δ double mutant as cells progress through mitosis. The occurence of sister chromatid separation (SCS) is indicated by the arrow. Horizontal scale bar: 2 min, vertical scale bar: 10 µm. **(C)** Cellular (left) and nuclear (right) concentration of separase-GFP in WT and *ufd2*Δ mutant as cells progress through mitosis. Mean (central line) and standard deviation (area) are shown (n = number of cells). The WT data shown here is the same as in Figure 3B. **(D)** Representative kymographs showing separase-GFP in wild-type (WT) and *ufd1-1* mutant as cells progress through mitosis. The occurence of sister chromatid separation (SCS) is indicated by the arrow. Horizontal scale bar: 2 min, vertical scale bar: 10 µm. The WT kymograph shown here is the same as in Figure 3B. **(E)** Cellular (left) and nuclear (right) concentration of separase-GFP in WT and *ufd1-1* mutant as cells progress through mitosis. The WT data shown here are the same as in Figure 3B. **(F)** Growth assay of *ufd1* mutants (*ufd1-1*, *ufd1-5* and *ufd1-3*), with or without moderate overexpression (8x)

### Figure S6
### Vijayakumari et al.

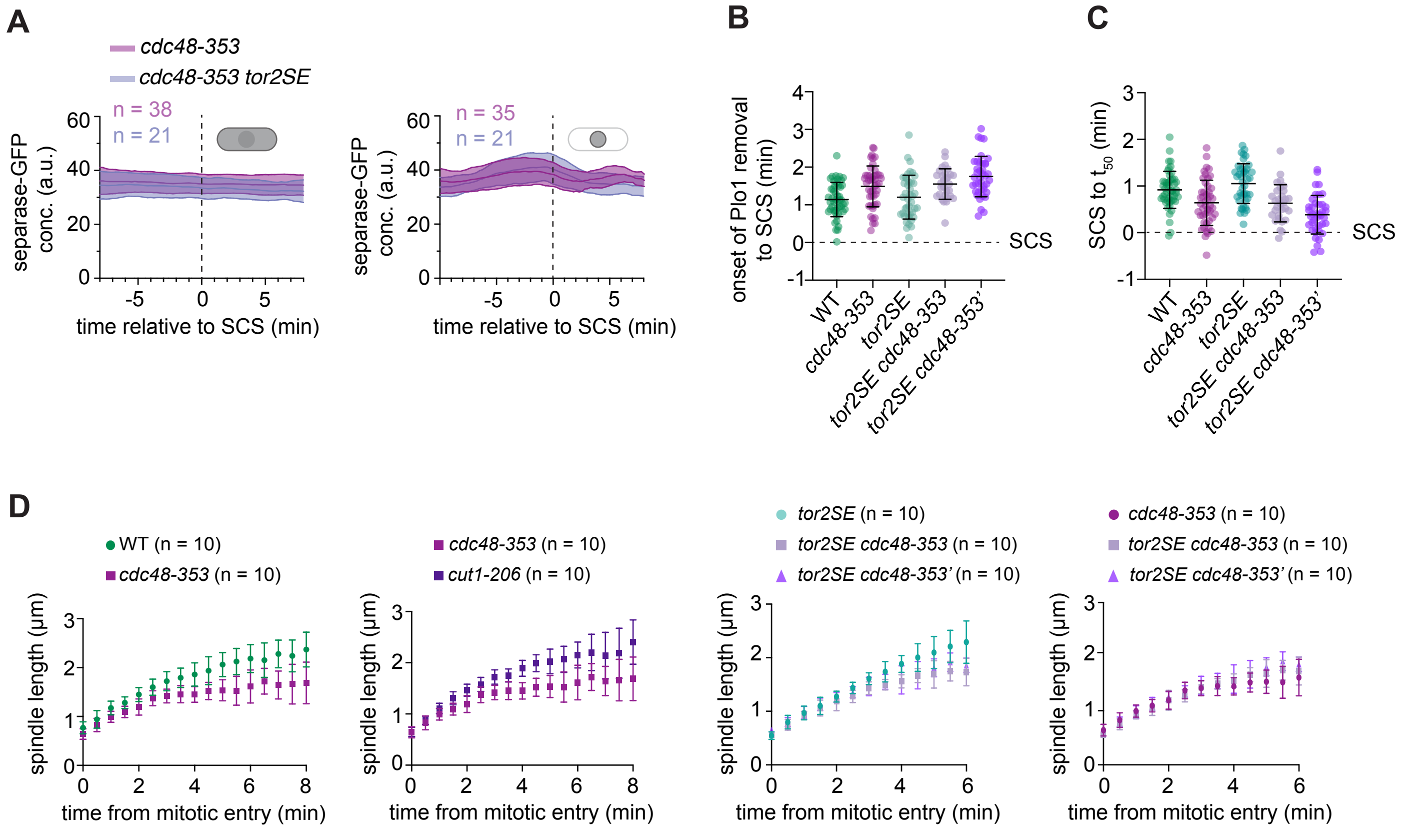

**Figure S6. Mutation of *tor2* neither rescues separase levels nor the short spindle elongation phenotype of *cdc48-353* mutant cells. (A)** Cellular (left) and nuclear (right) concentrations of separase-GFP in *cdc48-353* and *cdc48-353 tor2SE* double mutant during mitosis. Data are aligned to the time of sister chromatid separation (SCS), indicated by the vertical dashed line. Mean (central line) and standard deviation (area) are shown (n = number of cells). Data for the *cdc48-353* mutant is the same as that in Figure 3. **(B)** Time from onset of Plo1 removal to SCS in the indicated strains. Points are single cells, bars are mean and standard deviation. **(C)** Time from SCS to the point at which 50 % of Plo1 has been removed from the spindle pole bodies ($t_{50}$, mitotic exit) in the indicated strains. Points are single cells, bars are mean and standard deviation. **(D)** Measurements of spindle length during mitosis in WT and different mutants. Error bars are standard deviation. The same *cdc48-353* data are shown in the first two graphs, and the same double mutant data are shown in the third and fourth graph.

**Figure S7**
**Vijayakumari et al.**

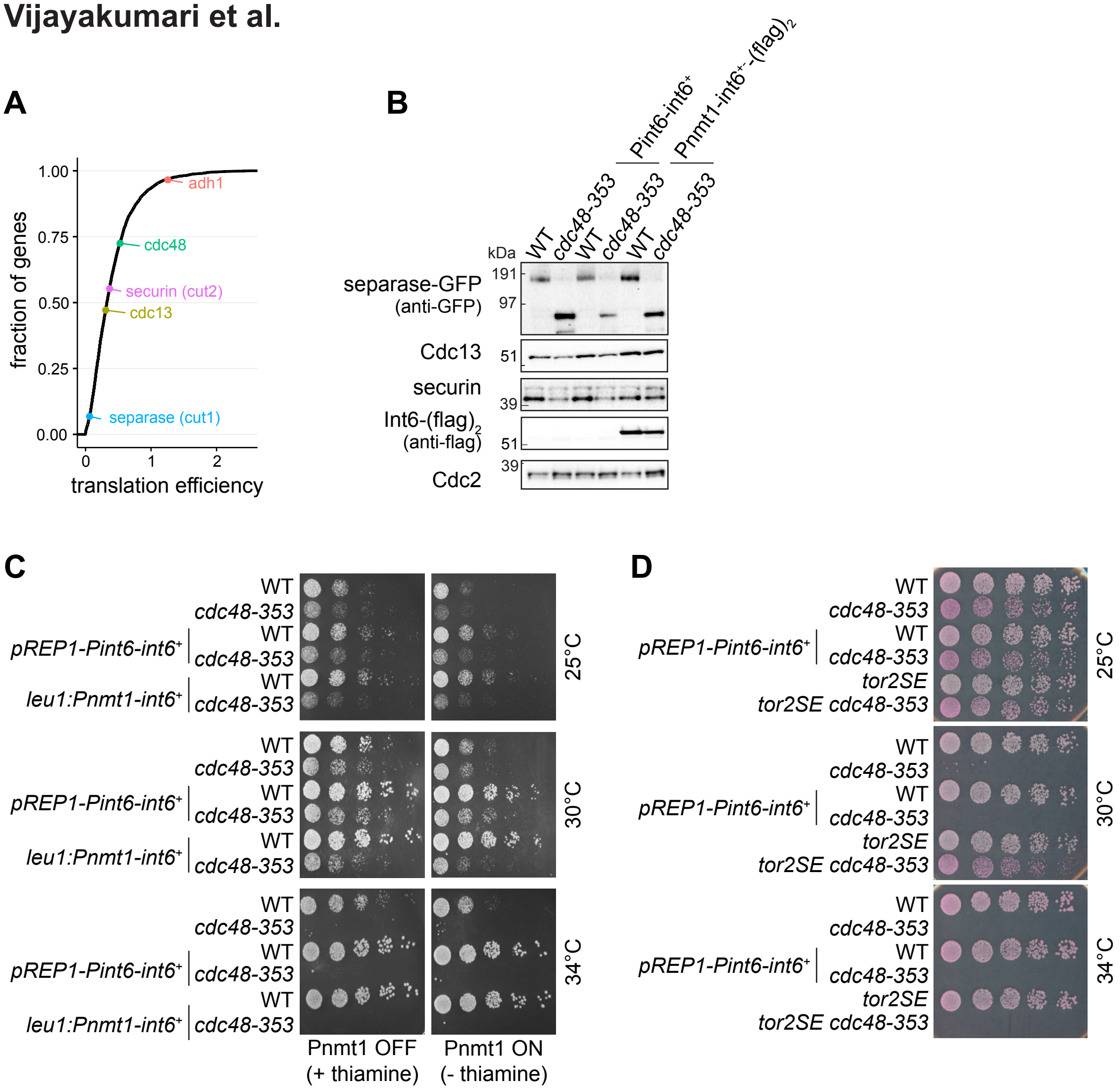

**Figure S7. Separase has low translation efficiency and Int6 overexpression does not rescue separase levels in the *cdc48-353* mutant. (A)** Cumulative frequency distribution showing the translation efficiency of *S. pombe* genes calculated as the ratio between previously reported raw counts of ribosome-protected RNA fragments and RNA-seq data (Rubio et al., 2020). In addition to separase (*cut1*) and *cdc48*, the *adh1* (alcohol dehydrogenase) gene is highlighted as an example of a highly expressed gene. **(B)** Immunoblot of the indicated proteins in WT and *cdc48-353* mutant with and without Int6 overexpression. **(C)** Growth of the indicated strains at different temperatures on minimal media (EMM) in the presence and absence of thaimine. **(E)** Growth of the indicated strains at different temperatures on rich medium containing Phloxine B, which stains dead cells.

### Table S1 - S. pombe strains

| | | |
|---|---|---|
| **Figure 1B, S1A-B, S3A** | | |
| SM318 | h+ | leu1 ade6-M216 cut1+-GFP<<kanR dh1L<<ura4+<<tetO Z<<natR<<Padh31-tetR-tdTomato |
| SW402 | h+ | leu1 cut1+-GFP<<kanR dh1L<<ura4+<<tetO Z<<natR<<Padh31-tetR-tdTomato cdc48-353 |
| **Figure 1C** | | |
| ST970 | h+ | plo1+-GFP<<kanR dh1L<<ura4+<<tetO Z<<natR<<Padh31-tetR-tdTomato |
| SW496 | h- | plo1+-GFP<<kanR dh1L<<ura4+<<tetO Z<<natR<<Padh31-tetR-tdTomato leu1 cdc48-353 |
| SW449 | n.d. | leu1 ade6-M216 plo1+-GFP<<kanR ura4-D18(?) dh1L<<ura4+<<tetO Z<<natR<<Padh31-tetR-tdTomato cut1-206 |
| **Figure 2A** | | |
| ST548 | h+ | cdc48-353 |
| ST224 | h- | leu1 ade6-M216 natNT2<<Padh1(#6)-cut2+ |
| **Figure 2B** | | |
| FY17242 | h- | leu1 cdc48-353 |
| **Figure 2C** | | |
| SW991 | h- | leu1 ade6-M216 cut1+-13myc<<kanR pREP81 |
| SW988 | h- | leu1 ade6-M216 cut1+-13myc<<kanR pREP81-Pnmt81-cut2+-GFP |
| SW989 | h- | leu1 ade6-M216 cut1+-13myc<<kanR pREP81-Pnmt81-cut2AIA-GFP |
| SW990 | h- | leu1 ade6-M216 cut1+-13myc<<kanR pREP81-Pnmt81-cut2N121-GFP |
| SX223 | h- | leu1 ade6-M216 cut1+-13myc<<kanR pREP81-Pnmt81-cut2 K104E, E129Y, D147G |
| SX224 | h- | leu1 ade6-M216 cut1+-13myc<<kanR pREP81-Pnmt81-cut2 L120F, E129N frameshift S167* |
| SW995 | h- | leu1 cdc48-353 cut1+-13myc<<kanR pREP81 |
| SW992 | h- | leu1 cdc48-353 cut1+-13myc<<kanR pREP81-Pnmt81-cut2+-GFP |
| SW993 | h- | leu1 cdc48-353 cut1+-13myc<<kanR pREP81-Pnmt81-cut2AIA-GFP |
| SW994 | h- | leu1 cdc48-353 cut1+-13myc<<kanR pREP81-Pnmt81-cut2N121-GFP |
| SX225 | h- | leu1 cdc48-353 cut1+-13myc<<kanR pREP81-Pnmt81-cut2 K104E, E129Y, D147G |
| SX226 | h- | leu1 cdc48-353 cut1+-13myc<<kanR pREP81-Pnmt81-cut2 L120F, E129N frameshift S167* |
| **Figure 2D** | | |
| SM336 | h- | leu1 ade6-M216 cut2+-GFP<<kanR cut1+-13myc<<kanR |
| SW991 | h- | leu1 ade6-M216 cut1+-13myc<<kanR pREP81 |
| SW988 | h- | leu1 ade6-M216 cut1+-13myc<<kanR pREP81-Pnmt81-cut2+-GFP |
| SW989 | h- | leu1 ade6-M216 cut1+-13myc<<kanR pREP81-Pnmt81-cut2AIA-GFP |
| SW990 | h- | leu1 ade6-M216 cut1+-13myc<<kanR pREP81-Pnmt81-cut2N121-GFP |
| SW995 | h- | leu1 cdc48-353 cut1+-13myc<<kanR pREP81 |
| SW992 | h- | leu1 cdc48-353 cut1+-13myc<<kanR pREP81-Pnmt81-cut2+-GFP |
| SW993 | h- | leu1 cdc48-353 cut1+-13myc<<kanR pREP81-Pnmt81-cut2AIA-GFP |
| SW994 | h- | leu1 cdc48-353 cut1+-13myc<<kanR pREP81-Pnmt81-cut2N121-GFP |
| **Figure 3A** | | |
| SL249 | h- | leu1 ade6-M216 cut2+-GFP<<kanR dh1L<<ura4+<<tetO Z<<natR<<Padh31-tetR-tdTomato |
| SW401p | h- | leu1 cut2+-GFP<<kanR dh1L<<ura4+<<tetO Z<<natR<<Padh31-tetR-tdTomato cdc48-353 |
| **Figure 3B** | | |
| SM318 | h+ | leu1 ade6-M216 cut1+-GFP<<kanR dh1L<<ura4+<<tetO Z<<natR<<Padh31-tetR-tdTomato |
| SW402 | h+ | leu1 cut1+-GFP<<kanR dh1L<<ura4+<<tetO Z<<natR<<Padh31-tetR-tdTomato cdc48-353 |
| **Figure 3C, S2B** | | |

| | | |
|---|---|---|
| SW402 | h+ | leu1 cut1+-GFP<<kanR dh1L<<ura4+<<tetO Z<<natR<<Padh31-tetR-tdTomato cdc48-353 |
| SW912 | h+ | leu1+<<pDUAL-Pcut1-cut1+-GFP cut1+-GFP<<kanR dh1L<<ura4+<<tetO Z<<natR<<Padh31-tetR-tdTomato cdc48-353 |
| SW910 | h+ | leu1+<<pDUAL-Pcut1-cut1+-GFP ade6-M216 cut1+-GFP<<kanR dh1L<<ura4+<<tetO Z<<natR<<Padh31-tetR-tdTomato |
| **Figure 4, S4** | | |
| SW922 | h+ | leu1 ade6-M216 cut1+-GFP<<kanR dh1L<<ura4+<<tetO Z<<natR<<Padh31-tetR-tdTomato pREP1 |
| SW918 | h+ | leu1 ade6-M216 cut1+-GFP<<kanR dh1L<<ura4+<<tetO Z<<natR<<Padh31-tetR-tdTomato pREP1-Pnmt1-(his)6-cdc48+ |
| SW919 | h+ | leu1 ade6-M216 cut1+-GFP<<kanR dh1L<<ura4+<<tetO Z<<natR<<Padh31-tetR-tdTomato pREP1-Pnmt1-(his)6-cdc48G338D |
| SW920 | h+ | leu1 ade6-M216 cut1+-GFP<<kanR dh1L<<ura4+<<tetO Z<<natR<<Padh31-tetR-tdTomato pREP1-Pnmt1-(his)6-cdc48E325A G338D |
| SW921 | h+ | leu1 ade6-M216 cut1+-GFP<<kanR dh1L<<ura4+<<tetO Z<<natR<<Padh31-tetR-tdTomato pREP1-Pnmt1-(his)6-cdc48M298Y |
| **Figure 5A-C, S5B,C** | | |
| SM318 | h+ | leu1 ade6-M216 cut1+-GFP<<kanR dh1L<<ura4+<<tetO Z<<natR<<Padh31-tetR-tdTomato |
| SW402 | h+ | leu1 cut1+-GFP<<kanR dh1L<<ura4+<<tetO Z<<natR<<Padh31-tetR-tdTomato cdc48-353 |
| SW936 | h+ | leu1 ade6-M216 cut1+-GFP<<kanR ufd1-5-myc:kanMX6 dh1L<<ura4+<<tetO Z<<natR<<Padh31-tetR-tdTomato |
| SW929 | n.d. | leu1 ade6-M216 cut1+-GFP<<kanR cdc48-353 ufd1-5-myc:kanMX6 |
| SW492 | h+ | leu1 ade6-M216 cut1+-GFP<<kanR ufd2::hph dh1L<<ura4+<<tetO Z<<natR<<Padh31-tetR-tdTomato |
| SW493 | h+ | leu1 cut1+-GFP<<kanR dh1L<<ura4+<<tetO Z<<natR<<Padh31-tetR-tdTomato cdc48-353 ufd2::hph |
| **Figure 5D-E** | | |
| SL249 | h- | leu1 ade6-M216 cut2+-GFP<<kanR dh1L<<ura4+<<tetO Z<<natR<<Padh31-tetR-tdTomato |
| SW484 | h- | leu1 ade6-M216 cut2+-GFP<<kanR dh1L<<ura4+<<tetO Z<<natR<<Padh31-tetR-tdTomato ufd2::hph |
| **Figure 5F** | | |
| SW481 | h- | ufd1-1-myc:kanMX6 leu1 ade6-M216 cut1+-GFP<<kanR dh1L<<ura4+<<tetO Z<<natR<<Padh31-tetR-tdTomato |
| SW402 | h+ | leu1 cut1+-GFP<<kanR dh1L<<ura4+<<tetO Z<<natR<<Padh31-tetR-tdTomato cdc48-353 |
| NBY3921 | h- | ufd1-5-myc:kanMX6 |
| **Figure 6** | | |
| SM318 | h+ | leu1 ade6-M216 cut1+-GFP<<kanR dh1L<<ura4+<<tetO Z<<natR<<Padh31-tetR-tdTomato |
| SW402 | h+ | leu1 cut1+-GFP<<kanR dh1L<<ura4+<<tetO Z<<natR<<Padh31-tetR-tdTomato cdc48-353 |
| SW940 | h- | tor2SE<<kanMX6 leu1 ade6-M216 cut1+-GFP<<kanR dh1L<<ura4+<<tetO Z<<natR<<Padh31-tetR-tdTomato |
| SW942 | h- | tor2SE<<kanMX6 leu1 cut1+-GFP<<kanR dh1L<<ura4+<<tetO Z<<natR<<Padh31-tetR-tdTomato cdc48-353 |
| SW941 | h- | fkh1::ura4+ leu1 ade6-M216 cut1+-GFP<<kanR dh1L<<ura4+<<tetO Z<<natR<<Padh31-tetR-tdTomato |
| SW944 | h- | fkh1::ura4+ leu1 cut1+-GFP<<kanR dh1L<<ura4+<<tetO Z<<natR<<Padh31-tetR-tdTomato cdc48-353 |

| | | |
|---|---|---|
| **Figure 7A-D** | | |
| SM318 | h+ | leu1 ade6-M216 cut1+-GFP<<kanR dh1L<<ura4+<<tetO Z<<natR<<Padh31-tetR-tdTomato |
| SW402 | h+ | leu1 cut1+-GFP<<kanR dh1L<<ura4+<<tetO Z<<natR<<Padh31-tetR-tdTomato cdc48-353 |
| SW951 | h+ | leu1 ade6-M216 cut1+-GFP<<kanR dh1L<<ura4+<<tetO Z<<natR<<Padh31-tetR-tdTomato atg1Δ::hph |
| SW952 | h- | cut1+-GFP<<kanR cdc48-353 atg1Δ::hph |
| SW949 | h+ | leu1 ade6-M216 cut1+-GFP<<kanR dh1L<<ura4+<<tetO Z<<natR<<Padh31-tetR-tdTomato pib1Δ::hph |
| SW950 | h+ | leu1 cut1+-GFP<<kanR dh1L<<ura4+<<tetO Z<<natR<<Padh31-tetR-tdTomato cdc48-353 pib1Δ::hph |
| SW967 | h+ | leu1 ade6-M216 cut1+-GFP<<kanR dh1L<<ura4+<<tetO Z<<natR<<Padh31-tetR-tdTomato psp3Δ::hph |
| SW968 | h+ | leu1 cut1+-GFP<<kanR dh1L<<ura4+<<tetO Z<<natR<<Padh31-tetR-tdTomato cdc48-353 psp3Δ::hph |
| SW961 | h+ | leu1 cut1+-GFP<<kanR dh1L<<ura4+<<tetO Z<<natR<<Padh31-tetR-tdTomato isp6Δ::hph |
| SW962 | h+ | leu1 cut1+-GFP<<kanR dh1L<<ura4+<<tetO Z<<natR<<Padh31-tetR-tdTomato cdc48-353 isp6Δ::hph |
| **Figure 7E** | | |
| SW997 | h- | leu1+<<Pnmt81-cut1+-GFP |
| SW998 | h- | leu1+<<Pnmt81-cut1+-GFP cdc48-353 |
| **Figure S1C-D** | | |
| ST957 | h+ | cdc13-internal-sfGFPcp dh1L<<ura4+<<tetO Z<<natR<<Padh31-tetR-tdTomato |
| SW404 | h+ | cdc13-internal-sfGFPcp dh1L<<ura4+<<tetO Z<<natR<<Padh31-tetR-tdTomato cdc48-353 |
| **Figure S2A,C,D** | | |
| SM318 | h+ | leu1 ade6-M216 cut1+-GFP<<kanR dh1L<<ura4+<<tetO Z<<natR<<Padh31-tetR-tdTomato |
| SW402 | h+ | leu1 cut1+-GFP<<kanR dh1L<<ura4+<<tetO Z<<natR<<Padh31-tetR-tdTomato cdc48-353 |
| SW913 | h+ | leu1+<<pDUAL-Pcut1-cut1+-13myc ade6-M216 cut1+-GFP<<kanR dh1L<<ura4+<<tetO Z<<natR<<Padh31-tetR-tdTomato |
| SW914 | h+ | leu1+<<pDUAL-Pcut1-cut1C1730A-13myc ade6-M216 cut1+-13myc dh1L<<ura4+<<tetO Z<<natR<<Padh31-tetR-tdTomato |
| SW915 | h+ | leu1+<<pDUAL-Pcut1-cut1+-13myc cut1+-GFP<<kanR dh1L<<ura4+<<tetO Z<<natR<<Padh31-tetR-tdTomato cdc48-353 |
| SW916 | h+ | leu1+<<pDUAL-Pcut1-cut1C1730A-13myc ade6-M216 cut1+-13myc dh1L<<ura4+<<tetO Z<<natR<<Padh31-tetR-tdTomato cdc48-353 |
| **Figure S2E** | | |
| SW476 | h- | cut1+-myc<<kanMX4 cdc25-22 tor2SE::kan ura4-D18 cdc48+-FRB-GFP<<natMX6 leu1:rpl13+-(FKB12)2 fkh1::ura4+ |
| **Figure S2F** | | |
| SW477 | h- | cut1+-GFP<<kanR tor2SE::kan ura4-D18 cdc48+-FRB<<natMX6 leu1+<<rpl13+-(FKB12)2 fkh1::ura4+ dh1L<<ura4+<<tetO Z<<natR<<Padh31-tetR-tdTomato |
| **Figure S3B** | | |
| ST577 | h- | leu1 cdc25-22 cut1+-13myc<<kanR plo1+-mCherry<<natR cut2+-GFP<<kanR |
| **Figure S5A** | | |
| SM318 | h+ | leu1 ade6-M216 cut1+-GFP<<kanR dh1L<<ura4+<<tetO Z<<natR<<Padh31-tetR-tdTomato |
| SW402 | h+ | leu1 cut1+-GFP<<kanR dh1L<<ura4+<<tetO Z<<natR<<Padh31-tetR-tdTomato cdc48-353 |
| SW718 | h+ | leu1 cut1+-GFP<<kanR plo1+-mCherry<<natR ubx4Δ::hphNT1 |

| | | |
|---|---|---|
| SW730 | ? | leu1 plo1+-mCherry<<natR cut1+-GFP<<kanR cdc48-353 ubx4Δ::hphNT1 |
| SW719 | h+ | leu1 cut1+-GFP<<kanR plo1+-mCherry<<natR otu1Δ::hphNT1 |
| SW720 | h+ | leu1 cdc48-353 cut1+-GFP<<kanR plo1+-mCherry<< natR otu1Δ::hphNT1 |
| SW721 | h+ | leu1 cut1+-GFP<<kanR plo1+-mCherry<<natR lub1Δ::hphNT1 |
| SW732 | h+ | leu1 plo1+-mCherry<<natR cut1+-GFP<<kanR cdc48-353 lub1Δ::hphNT1 |
| SW716 | h+ | ubx3Δ::natMX6 |
| SW717 | h+ | cdc48-353 ubx3Δ::natMX6 |
| **Figure S5D,E** | | |
| SM318 | h+ | leu1 ade6-M216 cut1+-GFP<<kanR dh1L<<ura4+<<tetO Z<<natR<<Padh31-tetR-tdTomato |
| SW481 | h- | ufd1-1-myc:kanMX6 leu1 ade6-M216 cut1+-GFP<<kanR dh1L<<ura4+<<tetO Z<<natR<<Padh31-tetR-tdTomato |
| **Figure S5F** | | |
| ST224 | h- | ade6-M216 leu1 natNT2<<Padh1(#6)-cut2+ |
| NBY3918 | h+ | ufd1-1-myc:kanMX6 |
| SW552 | h- | ade6-M216 leu1 natNT2<<Padh1(#6)-cut2+ ufd1-1 myc<<kanMX6 |
| NBY3921 | h- | ufd1-5-myc:kanMX6 |
| SW934 | h+ | ade6-M216 leu1 natNT2<<Padh1(#6)-cut2+ ufd1-5-myc:kanMX6 |
| NBY3919 | h+ | ufd1-3-myc:kanMX6 |
| SW553 | h- | ade6-M216 leu1natNT2<<Padh1(#6)-cut2+ ufd1-3 myc<<kanMX6 |
| **Figure S6A** | | |
| SW402 | h+ | leu1 cut1+-GFP<<kanR dh1L<<ura4+<<tetO Z<<natR<<Padh31-tetR-tdTomato cdc48-353 |
| SW942 | h- | tor2SE<<kanMX6 leu1 cut1+-GFP<<kanR dh1L<<ura4+<<tetO Z<<natR<<Padh31-tetR-tdTomato cdc48-353 |
| **Figure S6B-D** | | |
| ST970 | h+ | plo1+-GFP<<kanR dh1L<<ura4+<<tetO Z<<natR<<Padh31-tetR-tdTomato |
| SW996 | h- | plo1+-GFP<<kanR dh1L<<ura4+<<tetO Z<<natR<<Padh31-tetR-tdTomato leu1 cdc48-353 |
| SW986 | h- | tor2SE<<kanMX6 leu1 plo1+-GFP<<kanR dh1L<<ura4+<<tetO Z<<natR<<Padh31-tetR-tdTomato |
| SW987 | h- | tor2SE<<kanMX6 leu1 plo1+-GFP<<kanR dh1L<<ura4+<<tetO Z<<natR<<Padh31-tetR-tdTomato cdc48-353 |
| SW449 | n.d. | leu1 ade6-M216 plo1+-GFP<<kanR ura4-D18(?) dh1L<<ura4+<<tetO Z<<natR<<Padh31-tetR-tdTomato cut1-206 |
| **Figure S7B-D** | | |
| SM318 | h+ | leu1 ade6-M216 cut1+-GFP<<kanR dh1L<<ura4+<<tetO Z<<natR<<Padh31-tetR-tdTomato |
| SW402 | h+ | leu1 cut1+-GFP<<kanR dh1L<<ura4+<<tetO Z<<natR<<Padh31-tetR-tdTomato cdc48-353 |
| SX207 | h+ | leu1 pREP1-Pyin6-yin6+ ade6-M216 cut1+-GFP<<kanR dh1L<<ura4+<<tetO Z<<natR<<Padh31-tetR-tdTomato |
| SX208 | h+ | leu1 pREP1-Pyin6-yin6+ leu1 cut1+-GFP<<kanR dh1L<<ura4+<<tetO Z<<natR<<Padh31-tetR-tdTomato cdc48-353 |
| SX209 | h+ | leu1+<<Pnmt1-yin6+-(flag)2 ade6-M216 cut1+-GFP<<kanR dh1L<<ura4+<<tetO Z<<natR<<Padh31-tetR-tdTomato |
| SX210 | h+ | leu1+<<Pnmt1-yin6+-(flag)2 cut1+-GFP<<kanR dh1L<<ura4+<<tetO Z<<natR<<Padh31-tetR-tdTomato cdc48-353 |
| SW940 | h- | tor2SE<<kanMX6 leu1 ade6-M216 cut1+-GFP<<kanR dh1L<<ura4+<<tetO Z<<natR<<Padh31-tetR-tdTomato |
| SW942 | h- | tor2SE<<kanMX6 leu1 ade6-M216 cut1+-GFP<<kanR dh1L<<ura4+<<tetO Z<<natR<<Padh31-tetR-tdTomato cdc48-353 |
